# Genome-wide prediction of dominant and recessive neurodevelopmental disorder risk genes

**DOI:** 10.1101/2022.11.21.517436

**Authors:** Ryan S. Dhindsa, Blake Weido, Justin S. Dhindsa, Arya J. Shetty, Chloe Sands, Slavé Petrovski, Dimitrios Vitsios, Anthony W. Zoghbi

## Abstract

Despite great progress in the identification of neurodevelopmental disorder (NDD) risk genes, there are thousands that remain to be discovered. Computational tools that provide accurate gene-level predictions of NDD risk can significantly reduce the costs and time needed to prioritize and discover novel NDD risk genes. Here, we first demonstrate that machine learning models trained solely on single-cell RNA-sequencing data from the developing human cortex can robustly predict genes implicated in autism spectrum disorder (ASD), developmental and epileptic encephalopathy (DEE), and developmental delay (DD). Strikingly, we find differences in gene expression patterns of genes with monoallelic and biallelic inheritance patterns. We then integrate these expression data with 300 orthogonal features in a semi-supervised machine learning framework (mantis-ml) to train inheritance-specific models for ASD, DEE, and DD. The models have high predictive power (AUCs: 0.84 to 0.95) and top-ranked genes were up to two-fold (monoallelic models) and six-fold (biallelic models) more enriched for high-confidence NDD risk genes than genic intolerance metrics. Across all models, genes in the top decile of predicted risk genes were 60 to 130 times more likely to have publications strongly linking them to the phenotype of interest in PubMed compared to the bottom decile. Collectively, this work provides highly robust novel NDD risk gene predictions that can complement large-scale gene discovery efforts and underscores the importance of incorporating inheritance into gene risk prediction tools (https://nddgenes.com).

## Introduction

Neurodevelopmental disorders (NDDs), including autism spectrum disorder (ASD), developmental and epileptic encephalopathy (DEE), and developmental delay (DD), are highly heritable. Researchers have made great progress in identifying hundreds of genes associated with these disorders through sequencing studies of trios, families, and case-control cohorts ^1–7^. However, most patients with an NDD still do not receive a genetic diagnosis ^8^, in part because there are more NDD-associated genes to discover. In the case of ASD, only 190 of the estimated 1,000 risk genes have been confidently linked to disease ^9^, even as cohort sizes have grown to over 20,000 cases ^6^. Fully characterizing the genetic architecture of NDDs is crucial to making accurate molecular diagnoses, elucidating disease mechanisms, and developing targeted therapies but will likely require hundreds of thousands of additional sequenced cases^2^.

*In silico* approaches can help predict NDD risk genes and accelerate gene discovery. For example, we and others have shown that genes associated with severe early-onset disorders are under strong purifying selection and thus tend to be depleted of nonsynonymous variation in the general population ^10–14^. Genic intolerance metrics, which quantify the degree to which genes are intolerant to functional variation, have become a cornerstone in prioritizing NDD risk genes ^1,2,6,15–19^. However, not all intolerant genes are involved in NDDs, as any gene in which mutations reduce fecundity will be intolerant to variation (e.g., genes involved in fertility). Moreover, although population-level sequencing datasets continue to grow, intolerance metrics still suffer from a lack of power for smaller genes. Finally, perhaps the biggest current limitation is that although these scores can reliably detect purifying selection against variants with monoallelic/dominant inheritance patterns, they struggle to prioritize disease genes with biallelic/recessive modes of inheritance ^20–22^. Moreover, to our knowledge, there are no currently available disease-specific computational risk predictors for recessive disorders.

Other commonly used methods for predicting NDD risk genes rely on gene expression networks ^23,24^. However, most of these methods have been based on bulk RNA-sequencing data and thus do not account for potential cell type-specific expression patterns. Here we hypothesized that we could bolster NDD risk gene predictions by integrating genic intolerance, bulk- and single-cell RNA-sequencing data, and other orthogonal datasets in an inheritance-specific manner. First, we assess cell type-specific expression patterns for ASD, DEE, and DD genes stratified by inheritance pattern (i.e., monoallelic vs. biallelic). We then demonstrate that expression patterns alone can predict NDD risk genes but that these predictions significantly improve when used in combination with intolerance metrics. Finally, we use single-cell RNA-sequencing data, intolerance metrics, and hundreds of other gene-level annotations in a semi-supervised machine learning approach (mantis-ml) ^25^ to generate inheritance-specific risk gene predictions for ASD, DEE, and DD. Top risk gene predictions from these models show a striking enrichment for top genes from trio studies and large case-control analyses, expert-curated risk gene lists, and genes enriched for their related phenotype associations in published case reports and case series. We make the scores available through a public browser: https://nddgenes.com.

## Results

### Cell type enrichment of NDD risk genes

We examined the expression patterns of NDD risk genes using a recently published single-cell RNA-sequencing (scRNA-seq) atlas of the developing human cortex ^26^. This dataset contains 57,868 cells collected from four human fetal cortical samples spanning 8 weeks during mid-gestation, including post-conception week (PCW) 16, PCW20, PCW21, and PCW24 (**Fig. 1A, B**). There are 23 annotated cell types (**Fig. 1A**), including interneurons from the medial ganglionic eminence (MGE) and central ganglionic eminence (CGE), nine different clusters of cortical excitatory neurons (GluN), precursor cells like radial glia, and other non-neuronal cell types. One of the GluN clusters corresponds to the subplate, a transient cortical structure that contains some of the earliest formed neurons of the cortex (**Table S1**).

**Figure 1.**
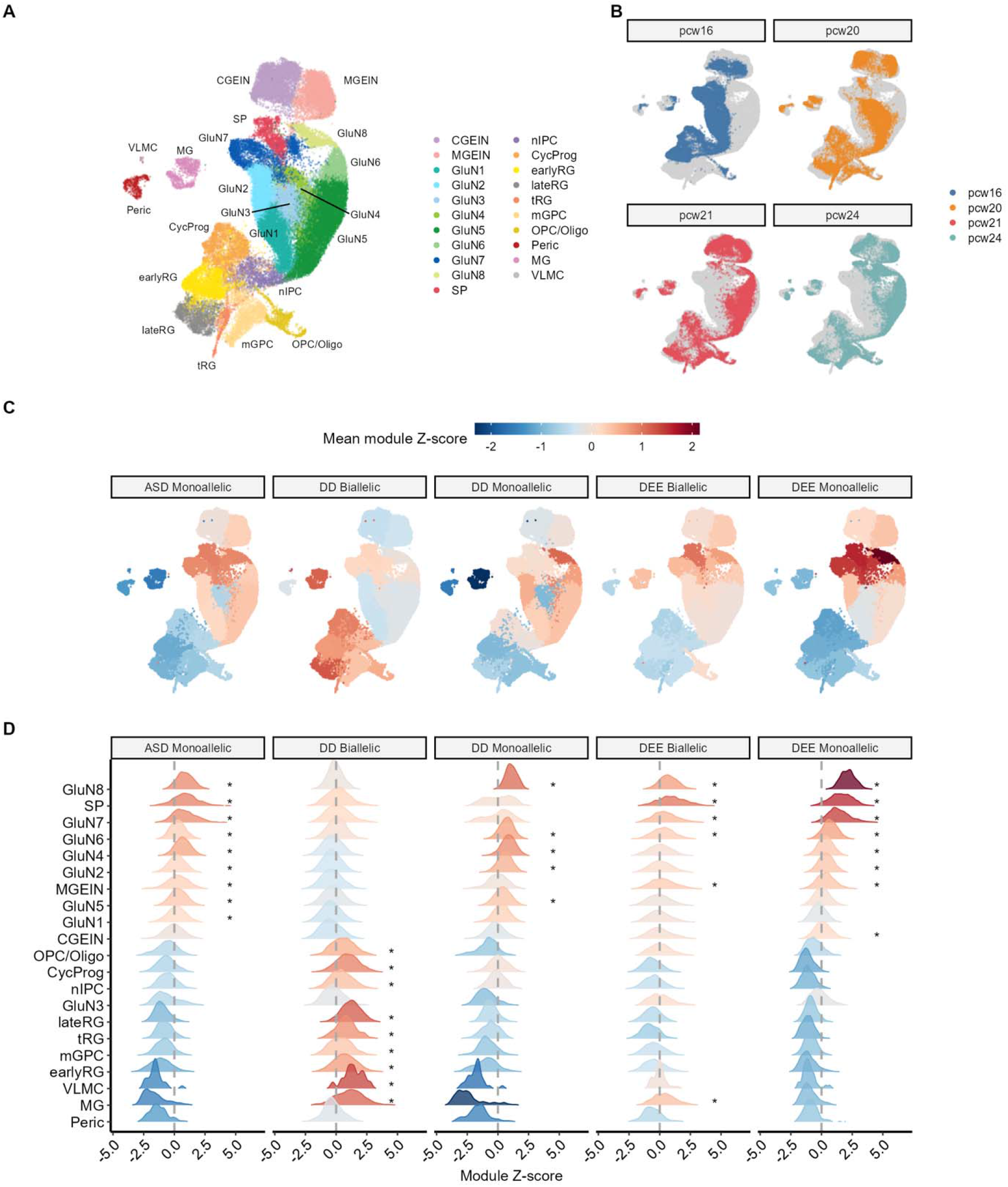
Cell type-specific expression patterns for NDD risk genes. **(A)** Uniform manifold approximation projection (UMAP) plot of the human fetal cortex from data generated by Trevino et al^26^. Cells colored by cell type. RG = radial glia; CycProg = cycling progenitors; tRG = truncated radial glia; mGPC = multipotent glial progenitor cell; OPC/Oligo = oligodendrocyte progenitor cell/oligodendrocyte; nIPC = neuronal intermediate progenitor cell; GluN = glutamatergic neuron; CGE IN = caudal ganglionic eminence interneuron; MGE IN = medial ganglionic eminence interneuron; EC = endothelial cell; MG, microglia; Peric. = Pericytes. **(B)** UMAP with cell types colored by age. **(C)** UMAP colored by module Z-scores for NDD gene sets. **(D)** Distribution of module Z-scores for each cell type. Asterisks indicate Bonferroni-corrected Mann-Whitney U p-values < 0.05.

To test whether NDD risk genes are preferentially expressed in any of these cell types, we carefully curated genes that have been implicated in ASD, DEE, and DD (Methods). We further annotated these genes as “monoallelic” or “biallelic” depending on the pattern of inheritance of pathogenic mutations in each gene (Methods). In total, we identified 190 monoallelic ASD genes, 94 monoallelic DEE genes, and 438 monoallelic DD genes. We also identified 17 biallelic ASD genes, 63 biallelic DEE genes, and 473 biallelic DD genes. We excluded biallelic ASD genes from downstream analyses due to the relatively small size of this gene set.

To determine whether each of these gene sets was more highly expressed in any fetal cortical cell type, we computed module Z-scores as previously described (see Methods).^27^ A positive Z-score indicates that the module of genes is expressed more highly in a particular cell than in the rest of the population. We calculated Mann-Whitney P-values for each cluster by randomly sampling 400 cells from the given cluster and comparing them to 400 random cells outside of that cluster. Monoallelic ASD genes were most significantly enriched in several glutamatergic neuron clusters, particularly those corresponding to more mature neurons (GluN4-8 and subplate) as well as MGE-derived interneurons (**Fig. 1D, Table S2**).

Monoallelic DEE genes showed a similar pattern as the ASD monoallelic genes, most strongly enriched in more mature GluN neurons (GluN 6-8), subplate neurons and MGE-derived interneurons. Biallelic DEE genes were enriched for GluN6, GluN7, GluN8, and SP neurons, but were not significantly enriched in MGE interneurons, though we note that we were less powered for this gene set given the smaller sample size compared to monoallelic genes. Monoallelic DD genes were also generally enriched in GluN neurons but were not significantly enriched in SP excitatory neurons or MGE interneurons.

Most interestingly, DD biallelic genes showed a strikingly different pattern from DD monoallelic genes and were preferentially expressed in more immature cell types and non-neuronal cells, such as oligodendrocyte precursor cells (OPCs), intermediate progenitor cells (IPCs), early and transitional radial glia (**Fig. 1D**). Altogether, these expression patterns support the notion that monoallelic ASD, DEE, and DD risk genes converge on similar cell types. However, while prior studies have suggested that DD genes are enriched for radial glia ^28^, we only observe a significant enrichment for biallelic DD genes in this cell type.

### Single-cell expression data bolsters NDD risk gene predictions

Motivated by their cell type-specific expression patterns, we hypothesized that we could leverage fetal single-cell RNA-sequencing data to predict NDD risk genes stratified by inheritance pattern. To this end, we trained random forest models using the scRNA-seq data for each of the NDD risk gene sets and compared their performance to models based on conventional intolerance metrics. We trained models for each disease gene list using the risk genes as the positively labeled set and a randomly selected set of genes as the negative set (1.5x the size of the risk gene list). We then compared model performance using five-fold cross-validation.

Random forest models trained purely on single-cell expression data could accurately predict NDD risk genes for each gene list (**Fig. 2A-E**). For monoallelic ASD, DD, and DEE, the random forest models achieved an area under the receiving operator curve (AUC) statistic of 0.85, 0.82, and 0.78, respectively. The monoallelic scRNA-seq models performed nearly as well as models trained with the loss-of-function observed/expected upper bound fraction (LOEUF) score, one of the most used loss-of-function intolerance metrics ^20^ (**Fig. 2A-C)**. Interestingly, for NDD risk genes with biallelic patterns of inheritance, scRNA-seq models outperformed models trained on any of the intolerance metrics (**Fig. 2D-E**). The expression profiles in the cell types with the highest module scores were among the most important features for each model (**Fig. S1**).

**Figure 2.**
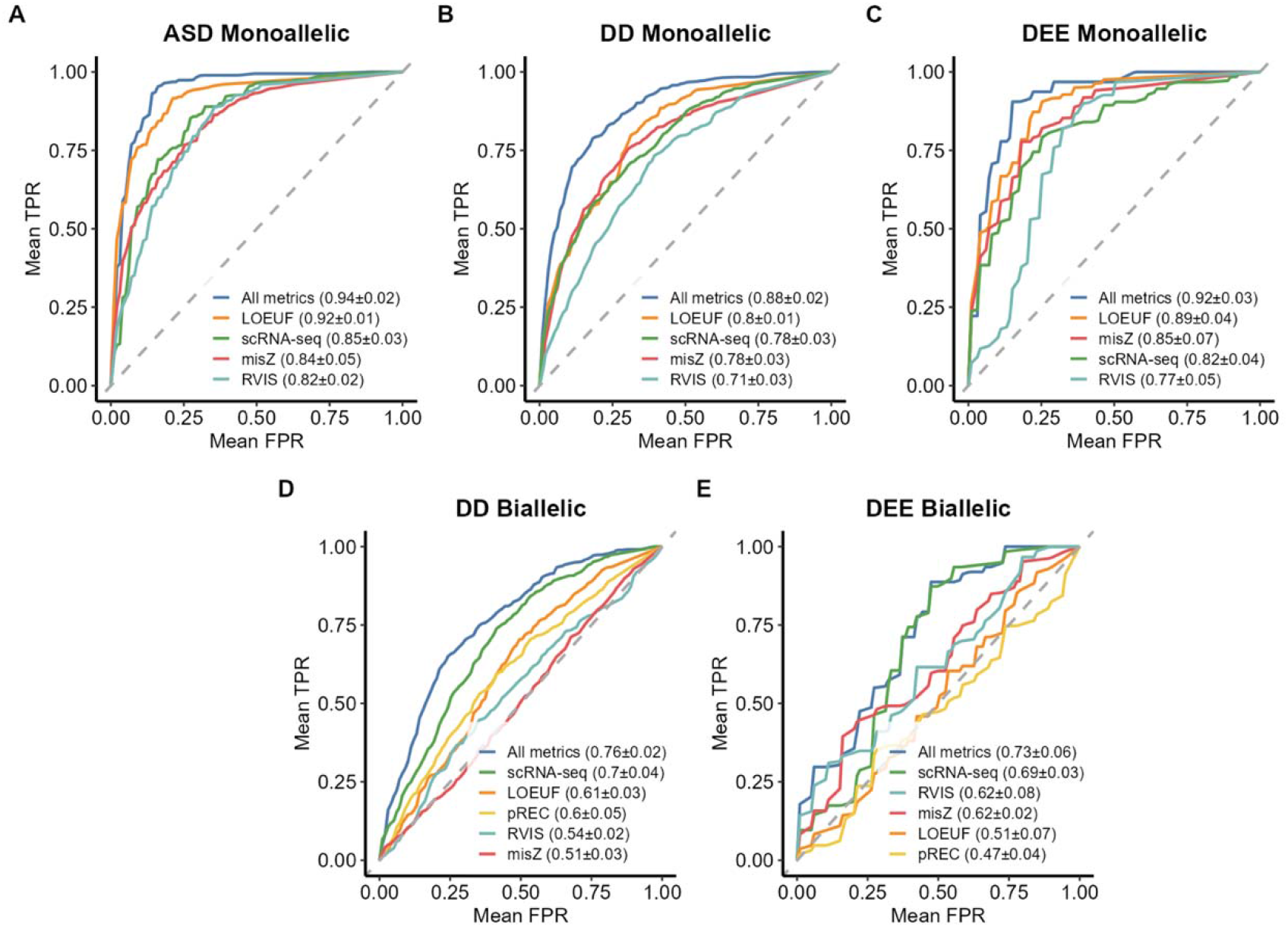
Random forest models including cortical single-cell expression data can predict NDD genes. Mean receiver operating characteristic (ROC) curves (from fivefold cross-validation) depcting the ability of random forest models trained with single-cell RNA-sequencing (scRNA-seq) data, intolerance metrics, or both (“All models”) for **(A)** monoallelic ASD genes, **(B)** monoallelic DD genes, **(C)** monoallelic DEE genes, **(D)** biallelic DD genes, and **(E)** biallelic DEE genes. TPR: True positive rate, FPR: False positive rate. The numbers in parentheses in each figure legend refer to the mean AUC and the standard deviation across the five folds.

We next investigated whether the expression-informed models were detecting information orthogonal to intolerance. To assess this, we built random forest models that incorporated both scRNA-seq data and intolerance metrics, including LOEUF, missense Z (misZ), and the residual variation intolerance score (RVIS) ^10^. We also included the probability of being intolerant to recessive variation (pREC) score, a measure of genic intolerance to biallelic loss-of-function variants, for the biallelic gene sets ^20^. The composite models consistently outperformed all individual models for each NDD subclass, regardless of the inheritance pattern (**Fig. 2**). Collectively, these results suggest that both scRNA-seq data and intolerance provide independent information in detecting NDD risk genes.

### Incorporation of scRNA-seq data in a semi-supervised machine learning model

One major challenge in generating genome-wide disease risk predictions is that although we have a set of known risk genes for each disease, we do not know which are definitively not associated with the disease (i.e., a true negative set). To address this, we previously introduced a stochastic semi-supervised machine learning approach called mantis-ml ^25^. Briefly, mantis-ml takes as input a list of seed genes (the positive set) and then trains machine learning models on random balanced datasets across the protein-coding exome. It then generates final gene rankings by averaging prediction probabilities across all the iterations. Mantis-ml includes several gene-level features, including several intolerance metrics, protein-protein interaction networks, and others.

Here, we made several advances to the mantis-ml framework. Foremost, we manually curated highly confident seed gene lists for ASD, DEE, and DD (**Table S3**). Given the differences in intolerance and expression profiles for monoallelic and biallelic gene sets, we trained inheritance-specific models. In addition, we include several new features, including scRNA-seq data and a new intolerance metric, gene variation intolerance rank (GeVIR), which was previously shown to be more sensitive for smaller genes ^29^. Finally, we introduce a gene ontology feature selection strategy, in which we perform enrichment analyses on the seed gene list to determine the gene ontologies to include as features in each model (Methods).

We trained separate mantis-ml models for monoallelic ASD, monoallelic DEE, monoallelic DD, biallelic DEE, and biallelic DD. The XGBoost models showed strong predictive power, with average AUCs of 0.95, 0.94, and 0.94 for monoallelic ASD, DEE, and DD and mean AUCs of 0.84 and 0.88 for biallelic DEE and DD, respectively (**Fig. 3B, Tables S4-8**Random forest models performed comparably, and we defaulted to the XGBoost-derived models for downstream analyses (**Table S9**). Using the Boruta algorithm, we computed feature importances for each XGBoost model (Methods; **Figures S2-4**). Constraint metrics were consistently among the top features for the monoallelic models, whereas expression data and protein-protein interaction data were relatively more important in the biallelic models.

**Figure 3.**
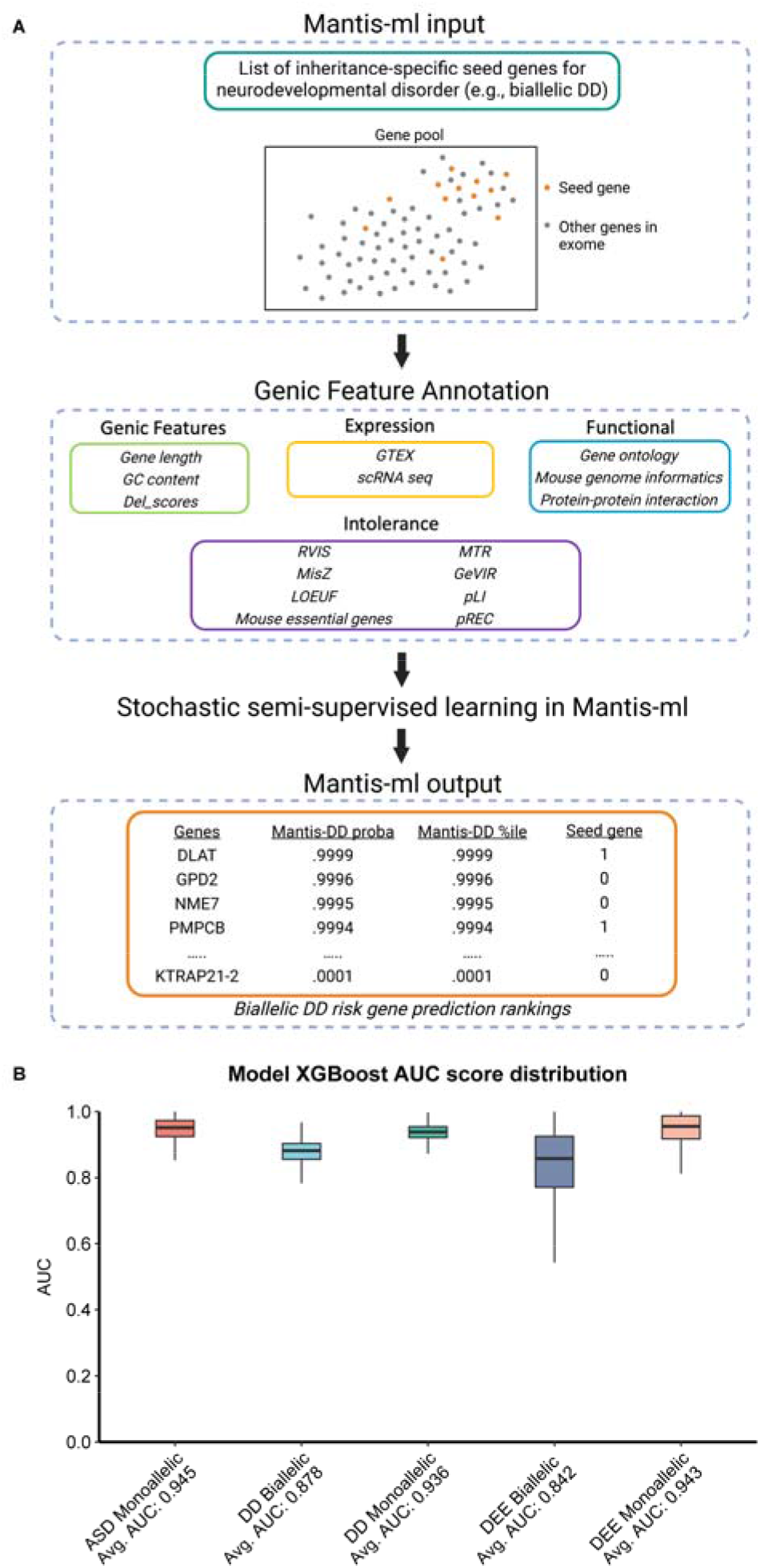
Mantis-ml XGBoost classifier performance across five neurodevelopmental disorder models. **(A)** Schematic of the mantis-ml framework, using biallelic developmental delay (DD) as an example seed gene list. **(B)** Score distribution of XGBoost area under the curves (AUC) across all five neurodevelopmental disorder risk mantis-ml models.

### Mantis-ml prioritizes top genes from rare variant association studies

We sought to evaluate mantis-ml’s ability to prioritize putative novel NDD risk genes using results from recent large-scale exome sequencing studies of ASD, DD, and epilepsy cohorts ^1,6^ (**Tables S11-13**). We thus tested whether top-ranked genes from each mantis-ml model were enriched for nominally significant genes (p<0.01) from these trio and case-control analyses.

Across all three dominant models, genes in the top 5^th^ percentile of mantis-ml were highly enriched for genes with nominal evidence of rare variant burden in ASD, DEE, and DD cases (p < 0.01) (ASD OR = 13, 95%CI: [10.7, 15.8], p = 3.5 × 10^−116^; DD OR = 25.3, 95%CI: [21.2, 30.2], p = 4.9 × 10^−244^; DEE OR = 16.8, 95%CI: [6.4, 44.2], p = 1.0 × 10^−8^). These enrichments remained highly significant even after removing seed genes from the evaluation (ASD OR = 8.5, 95%CI: [6.7, 10.8], p = 1.2 × 10^−52^; DD OR = 13.4, 95%CI: [10.6, 16.8], p = 4.7 × 10^−81^; DEE OR = 12.3, 95%CI: [3.5, 40.5], p = 7.4 × 10^−05^).

We then performed the same enrichment tests using LOEUF instead of mantis-ml. Of note, the ASD and DD exome studies used LOEUF as a gene weight in their burden tests ^6,7^, meaning that top-ranked hits would be skewed for more LOF-intolerant genes. Despite this, nominally significant (p<0.01) genes from the ASD, DD, and DEE studies were less strongly enriched for genes within the top 5% of LOEUF than with mantis-ml (Fig. 4). For example, in the DD study (the best powered of the three studies), top mantis-ml genes (with seed genes removed) had an odds ratio of 13.4 (95%CI: [10.6, 16.8]; p = 4.7 × 10^−81^) compared to an odds ratio of 8.2 (95%CI: [6.4, 10.4]; p =2.0 × 10^−50^) for top-ranked LOEUF genes. This suggests that mantis-ml has nearly twice the ability to prioritize potential novel DD risk genes when compared to LOEUF and represents a significant improvement over the current standard in the field. Finally, we compared the performance of these monoallelic-specific models to mantis-ml models that were trained on seed gene lists that were not stratified by inheritance. The inheritance-informed monoallelic models substantially outperformed the inheritance-agnostic models for both DD and DEE, with 1.6- and 2.8-times larger point estimates, respectively (**Fig. S5, Table S14 and S15**).

**Figure 4.**
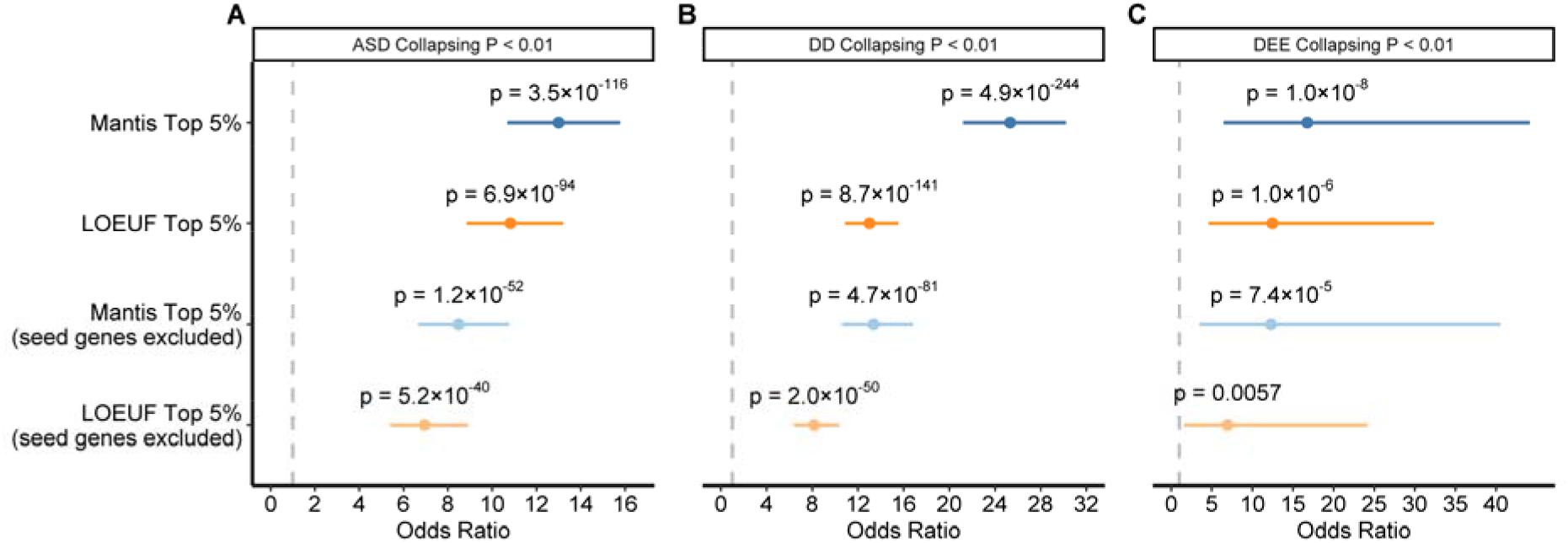
Enrichment of mantis-ml predictions among top genes from rare variant gene-level association studies. **(A-C)** The enrichment of top mantis-ml predictions (≥ 95^th^ percentile) and LOF-intolerant genes (measured via LOEUF) among nominally significant (p<0.01) genes in prior gene-level association studies for ASD (n cases **=** 20,627), DD (n cases = 31,058), and DEE (n cases = 1,021), respectively. Mantis-ml models for each figure represent the monoallelic model for each respective NDD. Error bars represent 95% confidence intervals. P-values calculated via two-tailed Fisher’s exact test. Bonferroni-corrected p-value threshold = 0.004 for an alpha of 0.05.

### Mantis-ml risk predictions align with the degree of confidence in clinically curated gene lists

We next tested how well the mantis-ml predictions correlated with manually curated NDD risk gene lists, including those from the Simons Foundation Autism Research Initiative (SFARI) database of ASD genes ^30^ and the DECIPHER Developmental Disorder Genotype–Phenotype Database (DDG2P) of DD genes ^31^ (**Table S16 and S17**). In both resources, each gene receives a score reflecting the strength of evidence in the published literature of a gene’s role in the disease. SFARI ranks genes by Tier, in which Tier 1 includes “high confidence” genes (n = 204), Tier 2 includes “strong candidate” genes (n = 208), and Tier 3 includes genes with “suggestive evidence” (n = 493). The DDG2P resource includes “Definitive” (n = 218), “Strong” (n = 156), and “Limited” (n = 63) categories for monoallelic risk genes and “Definitive” (n = 452), “Strong” (n = 202), and “Limited” (n = 98) for biallelic risk genes. Genes from SFARI’s Tier 1 and DDG2P’s “Definitive” category and a subset of monoallelic genes from DDG2P’s “Strong” category (n = 55) were used as seed genes for our models, providing an opportunity to test mantis-ml’s performance on the remaining gene lists (e.g., Tier 2/3 and “Strong”/”Limited”), which mostly consist of genes that have emerged from smaller trio- and family-based sequencing studies and functional validation.

We found that the distribution of mantis-ml percentiles correlated with the levels of evidentiary support and expert curation for both ASD and DD (**Fig. 5**). As expected, the seed genes had the highest mantis-ml percentiles (**Fig. 5A-C**), which were significantly higher than the remaining genes in the exome (monoallelic ASD MWU p = 3.2 × 10^−93^, monoallelic DD MWU p = 1.7 × 10^−110^, biallelic DD MWU p = 2.9 × 10^−176^). The percentile ranks of Tier 2 SFARI genes and DD2GP “Strong” genes were on average lower than seed genes but significantly higher than the rest of the exome (**Fig. 5A-C)**. Likewise, the percentiles of Tier 3 SFARI genes and “Limited” DD genes were still significantly higher than the remaining genes in the exome, but not as enriched as the higher confidence gene sets (**Fig. 5A-C)**.

**Figure 5.**
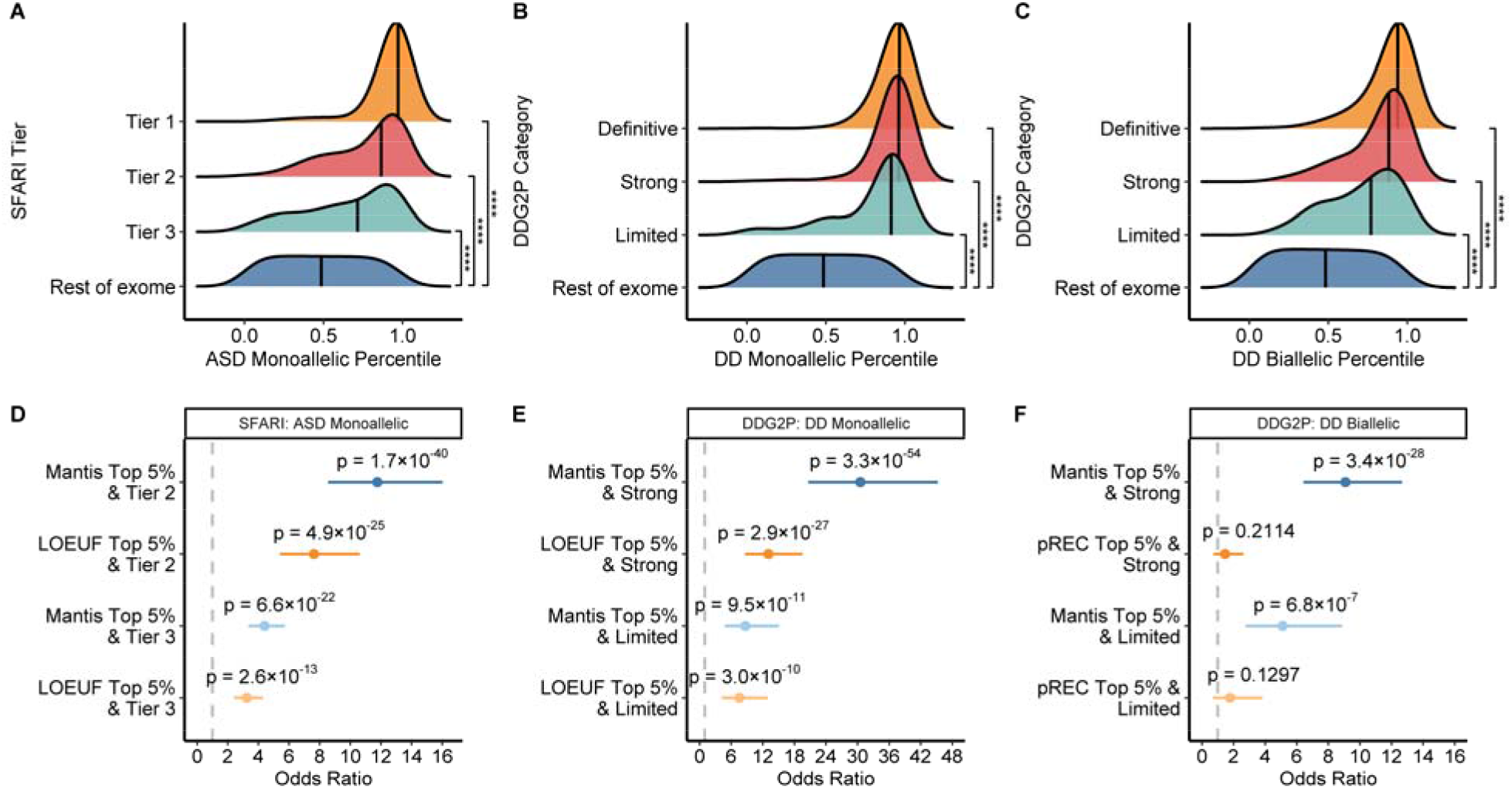
Mantis-ml performance across rare variant association studies and clinically curated gene lists. (**A-C**) The distribution of mantis-ml risk percentiles among clinically curated gene lists from SFARI and DDG2P compared to the rest of the exome, respectively. ASD and DD seed genes were comprised of inheritance-specific SFARI Tier 1 genes and DDG2P “Definitive” genes, respectively. P-values were calculated via the Mann-Whitney U test. **** indicates Bonferroni-corrected p < 1×10^−14^. **(D)** Forest plots comparing the magnitude of SFARI Tier 2 and Tier 3 gene enrichment in top 5^th^ percentile monoallelic ASD mantis-ml predictions and LOEUF rankings. (**E)** Enrichment of DDG2P “Strong” and “Limited” gene categories in the top 5^th^ percentile of monoallelic DD mantis-ml predictions and LOEUF rankings. (**F**) DDG2P gene enrichment in top 5^th^ percentile of biallelic DD mantis-ml predictions and pREC scores. ASD: autism spectrum disorder, DD: developmental delay, SFARI: Simons Foundation Autism Research Initiative, DDG2P: Developmental Disorder Genotype–Phenotype Database.

We next compared the enrichment of mantis-ml predictions to intolerance metrics for these expert-curated gene lists (**Table S18 and S19**). Consistent with the collapsing analysis enrichment tests, Tier 2 and Tier 3 SFARI genes were more strongly enriched for top-ranked (top 5^th^ percentile) mantis-ml monoallelic ASD genes than the top 5^th^ percentile of LOEUF genes (Tier 2: OR = 11.8, 95% CI: [8.6, 16.0], p = 1.7 × 10^−40^, versus OR = 7.6, 95% CI: [5.4, 10.6], p = 4.9 × 10^−25^; Tier 3: OR = 4.4, 95% CI: [3.3, 5.7], p = 6.6 × 10^−22^ versus OR = 3.2, 95% CI: [2.4, 4.3], p = 2.6 × 10^−13^). Likewise, mantis-ml monoallelic DD predictions were more strongly enriched among “Strong” and “Limited” monoallelic DD2GP genes (“Strong”: OR = 30.5, 95% CI: [16.5, 38.2], p = 6.1 × 10^−41^ versus OR = 13.1, 95% CI: [8.4, 20.2], p = 3.5 × 10^−24^; “Limited”: OR = 9.9, 95% CI: [5.2, 17.8], p = 2.4 × 10^−10^, versus OR = 8.1, 95% CI: [4.2, 14.6], p = 3.6 × 10^−9^). Although the confidence intervals of these enrichments overlapped, the consistently higher point estimates for the top mantis-ml genes suggest that these predictions have a stronger discriminatory ability than LOEUF alone.

We observed an even more dramatic difference in enrichments among biallelic DD genes. We compared our biallelic DD mantis-ml predictions to the pREC intolerance score,^20^ which aims to capture the probability a gene is intolerant to homozygous loss-of-function. The top mantis-ml genes (top 5^th^ percentile) were strongly enriched for both “Strong” and “Limited” biallelic DDG2P genes. On the other hand, neither of these gene lists was significantly enriched for genes in the top 5^th^ percentile of pREC (OR = 1.5, 95 CI: [0.7, 2.6], p = 0.2 and OR = 1.8, 95% CI: [0.7, 3.8], p = 0.1, respectively). We compared the performance of the inheritance-stratified models versus the inheritance-agnostic models. For monoallelic DD, the odds ratios of the monoallelic DD models were 1.3-times and 1.08-times larger than for the inheritance-agnostic DD model for the “Strong” and “Limited” gene lists, respectively (**Fig. S6**). Likewise, the odds ratios for the DD-specific models were 1.9- and 1.2-times larger for biallelic “Strong” and “Limited” DD genes, respectively (**Fig. S6**). These results suggest that the biallelic DD mantis-ml model could substantially help in the discovery of biallelic risk genes, whose discovery typically requires access to consanguineous populations or very large sample sizes (**Table S20**).

### Mantis-ml flags genes in clinically curated databases with limited evidentiary support

The SFARI and DECIPHER DDG2P databases provide clinicians and researchers with broad categories of confidence for a gene’s relevance to ASD and DD, respectively. Genes within each category are considered to have the same level of evidentiary support. We sought to evaluate mantis-ml’s ability to provide a more nuanced and quantitative measure of NDD risk within these broad, manually curated categories. For each evidentiary category (i.e., Tier 2/3 in SFARI and “Strong”/”Limited” in DDG2P), we first separated genes into high (≥ 90^th^ percentile) and low (<50^th^ percentile) mantis-ml risk prediction groups. We removed any monoallelic DD seed genes that were included in the “Strong”/”Limited” categories. There are no ASD seed genes in Tier 2/3. We then used two orthogonal validations to corroborate mantis-ml’s predictions for each gene: publications linking a given gene to either ASD or DD and statistical support (p values) from the largest ASD/DD sequencing study to date^6^. To maximize statistical power, we combined genes from SFARI Tiers 2 and 3 for ASD and DDG2P “Strong”/”Limited” for DD, respectively.

We systematically assessed whether mantis-ml predictions correlated with the degree of literature support for each gene in SFARI/DDG2P using Automatic Mendelian Literature Evaluation (AMELIE) ^32^ (**Fig. S7**). AMELIE is a natural language processing tool that searches all of Pubmed for manuscripts that link genes to a phenotype of interest. Importantly, AMELIE can also detect whether there is language in each article that suggests a specific pattern of inheritance, which allowed us to search gene-phenotype relationships in an inheritance-specific manner. For SFARI Tier 2 and 3 genes, we found that 49.3% (108 out of 219) of high mantis-ml risk genes had ≥ 1 publications linking them to ASD compared to 13.4% (24 out of 179) of low mantis-ml risk genes (OR 6.3, 95%CI: [3.7, 10.9], p = 9.9×10^−15^). For the DDG2P “Strong”/”Limited” categories, 69.7% (154 out of 221) of high mantis-ml risk genes had ≥ 1 publication linking them to DD compared to 53.4% (31 out of 58) of low mantis-ml risk genes (OR 2.2, 95%CI: [1.2, 4.0], p = 0.02).

We next assessed statistical human genetics evidence support from the largest and most recent sequencing study of ASD and DD^6^ (**Fig. S8**). For SFARI Tier 2 and 3 genes, we found that 25.1% (52 out of 207) of high-mantis-ml risk genes had nominally significant p-values <0.01 compared to 0% (0 out of 173) of low mantis-ml risk genes (OR Inf, 95%CI: [4.3, Inf], p = 3×10^−6^). Similarly, for monoallelic DECIPHER “Strong”/”Limited” categories, 44.6% (45 out of 101) of high mantis-ml risk genes vs. 0% (0 out of 15) of low mantis-ml risk genes had p-values <0.01 (OR Inf, 95%CI: [2.7, Inf], p = 4×10^−4^). Of note, X chromosome genes were not included in the p-value analysis as they were not analyzed in the Fu et al. study.

These data demonstrate mantis-ml’s ability to flag likely false positive genes that are included in clinically curated databases such as SFARI and DECIPHER. For example, *CDH15* (mantis 46^th^ percentile) is a DDG2P “Limited” gene and currently has an active gene-phenotype listing in Online Mendelian Inheritance in Man (OMIM).^33^ However, the evidence for this association is supported only by one publication from 2008 which lists three missense variants that were purported to be associated with severe intellectual disability.^34^ A curation of these variants reveals that two have been reclassified as Benign in ClinVar and the third is present in 18 individuals in the gnomAD database, which is inconsistent with a pathogenic variant for severe intellectual disability. Similarly, *CD96* (mantis 3^rd^ percentile) is a DDG2P “Limited” gene with an active gene-phenotype listing for C Syndrome in OMIM. This association is only supported by one manuscript from 2007 which identified a translocation breakpoint in *CD96* in a patient with C syndrome and a missense mutation (Thr280Met) in *CD96* in a patient with Bohring-Opitz Syndrome.^35^ However, subsequent papers have largely refuted this association including a balanced translocation disrupting *CD96* without symptoms of C syndrome,^36^ negative mutation screening of *CD96* in C syndrome patients,^37^ phenotypically normal *Cd96-/-* mice,^38^ and the presence of the Thr280Met missense variant in six individuals in the non-neurologic subset of gnomAD. These are only two of many examples of genes flagged by mantis-ml as being unlikely to be causal for NDDs.

Manually curating databases such as SFARI, DECIPHER, and OMIM is a time-consuming process and prone to false positives given the vast amounts of literature and human genetics evidence that needs to be reviewed for thousands of genes. Given that clinicians often look to these databases when assessing the evidence for a gene’s involvement in a disease, it is critical to ensure that the genes included in these databases are of high quality. Our data show that mantis-ml can provide an automated, immediate, and inheritance-specific assessment of the evidence for each gene’s risk for NDDs that can aid clinicians and researchers who manually curate these databases.

### Mantis-ml predicts gene-phenotype relationships in published literature

Before emerging as significant in large-scale sequencing studies, genes are often initially implicated in disease through case reports with supporting functional work, case series, or family-based studies. Thus, we sought to evaluate the relationship between a gene’s predicted mantis-ml risk percentile and the number of publications linked to the phenotype of interest. We used AMELIE to identify the number of publications linking each gene in the genome to our three phenotypes of interest (ASD, DD, DEE) in an inheritance-specific manner.

For each mantis-ml model, we removed seed genes and binned the remaining genes into predicted mantis-ml risk deciles. Across all five models, the top mantis-ml deciles were significantly more enriched for genes with at least one publication linking the gene to the phenotype of interest when compared to the rest of the genes in the genome (**Fig. 6, Fig. S9 and Table S21**). The enrichments were even stronger when we considered the top first percentile (**Fig. 6**). There was a stepwise decrease in the strength of enrichment for each successive decile. We imposed a more stringent AMELIE cutoff in which we tested the enrichment of genes with at least five phenotype-matching PubMed records (the maximum allowed by AMELIE) and observed even stronger enrichments among the top deciles and first percentile for each model (**Fig. 6, Fig. S10**).

**Figure 6.**
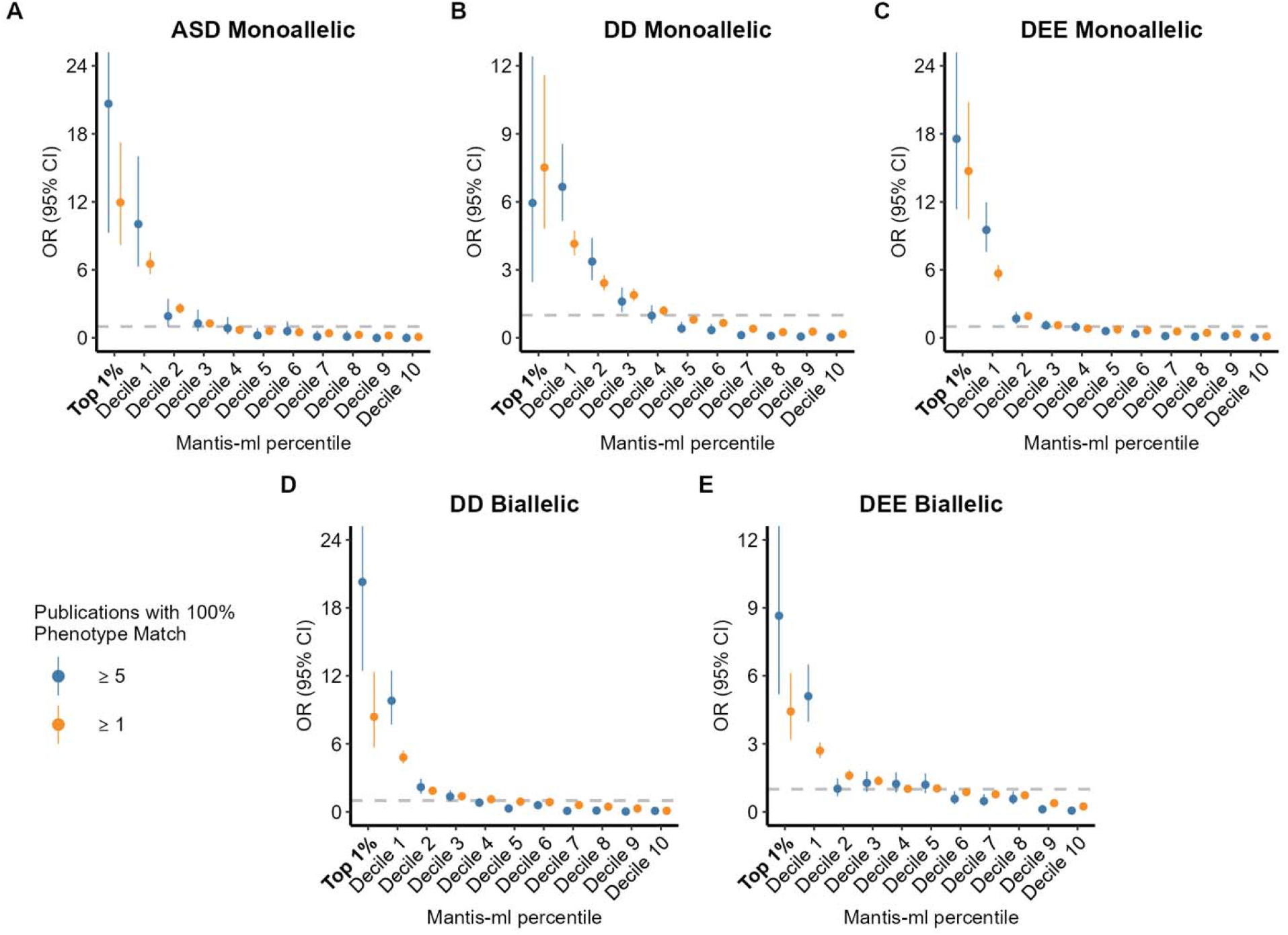
Enrichment of genes with 100% phenotype match from the published literature stratified by mantis-ml decile. For each model, we used AMELIE to generate gene-phenotype match scores from the literature for all genes in an inheritance specific manner. AMELIE gene-phenotype match scores range from 0 to 100%. We limited our analysis to 100% gene-phenotype matches from the literature based on the following phenotypes: HP:0000729 (autistic behavior) for ASD, HP:0012759 (neurodevelopmental abnormality) for DD, and HP:0001250 (seizures) for DEE. We then plotted the enrichment of gene-phenotype matches stratified by mantis-ml prediction deciles (and top 1st percentile) with ≥ 1 matching publications in orange and ≥ 5 in blue. P-values for these comparisons are available in Tables S21 and S22.

These results further support the role of mantis-ml in discriminating putative new NDD risk genes. For example, in the mantis-ml biallelic DD model, 8.8% (138/1575) of genes in the top decile have at least five publications linking them to biallelic DD in AMELIE vs. 0.2% (3/1861) of genes in the last decile (OR = 59.4, 95% CI: [19.8, 289.8], p = 5.0×10^−44^). Strikingly, 23.7% of genes in the top first percentile of risk have five or more publications linked to biallelic DD. Thus, these top percentile genes are more than 190 times more likely than genes in the bottom decile to have a high confidence association with biallelic DD in the literature (OR = 190.3 95%CI: [56.9, 1019.8], *p* = 5.4×10^−32^). The powerful discriminatory ability of mantis-ml between the top and bottom decile of predictions is consistent across all five disease models and inheritance patterns (ASD monoallelic OR Inf, 95%CI: [11.6, Inf], p = 1.1×10^−13^, DEE monoallelic OR 91.3, 95%CI: [24.8, 752.8], *p* = 2.3×10^−48^, DEE biallelic OR 59.1, 95%CI: [15.9, 494.0], p = 1.8×10^−31^, DD monoallelic OR 138.1, 95%CI: [24.2, 5338.7], p = 1.9×10^−36^) (**Fig. 6, Table S22**).

Lastly, we used AMELIE (≥ 5 publications) to compare the performance of mantis-ml models trained using inheritance-specific versus inheritance-agnostic seed gene lists for DD and DEE **(Table S23**). The enrichment of top decile mantis-ml hits was 1.4-, 1.3-, 2.5-, and 4.6-times greater for monoallelic DD, monoallelic DEE, biallelic DD, and biallelic DEE, respectively, for the inheritance-informed models (**Fig. S11**). For the biallelic models, these enrichments were even more striking in the top percentile of mantis-ml risk, with odds ratios that were 9.7-times and 21.5-times higher for biallelic DD and biallelic DEE, respectively (**Fig. S11**).

## Discussion

While there has been great progress in identifying hundreds of genes associated with neurodevelopmental disorders, there remain thousands of additional risk genes to be identified. Sequencing studies will require hundreds of thousands of additional participants to fully resolve the genetic architecture of neurodevelopmental disorders^2^. Here, we used the mantis-ml semi-supervised machine-learning framework to provide dominant and recessive gene risk predictions across the spectrum of neurodevelopmental disorders. We conducted multiple orthogonal validations of mantis-ml that demonstrate its ability to prioritize both monoallelic and biallelic risk genes for neurodevelopmental disorders.

Our results suggest that the monoallelic ASD, DD, and DEE models outperform intolerance metrics alone in prioritizing the top results from NDD rare variant association studies. While intolerance metrics such as LOEUF, RVIS, and others have proven extremely useful in prioritizing risk genes, they are not specific to any disease. Mantis-ml leverages multiple measures of genic intolerance, bulk, and single-cell RNAseq data, protein-protein interaction networks, and gene ontology annotations tailored to the specific disorder and inheritance pattern of interest. We also showed that mantis-ml predictions aligned with experts’ degree of confidence in risk genes included in curated gene lists available through SFARI and DDG2P. However, mantis-ml also flagged several genes in the SFARI and DDG2P databases as having a low likelihood of being risk genes for ASD or DD. We showed that SFARI/DDG2P genes with low mantis-ml risk percentiles (<50^th^ percentile) for ASD/DD have significantly fewer publications and weaker human genetics supporting evidence from the largest ASD/DD sequencing studies, suggesting that they are unlikely to be true risk genes. These results suggest that mantis-ml predictions can help geneticists further prioritize disease genes in clinically curated lists and that one should reconsider the evidence for those with very low mantis-ml predictions. To this point, *KATNAL2*, currently a Tier 1 SFARI gene (the highest confidence), was predicted by the monoallelic ASD model to have only a 2.9% chance of being an ASD risk gene, despite being used as a seed gene in our original analysis. Indeed, a recent re-curation of the evidence for *KATNAL2* as a risk gene suggests that it is unlikely to contribute to autism risk through haploinsufficiency and it is no longer statistically significant in the most recent and largest ASD sequencing study ^6,39^.

We foresee several clinical applications for mantis-ml. First, mantis-ml can be used in conjunction with genomic or functional evidence to accelerate gene discovery. For example, mantis-ml can provide orthogonal evidence to prioritize genes with strong human genetics evidence that do not yet meet genome-wide significance in association studies. Second, we have shown that mantis-ml can also substantially improve the reliability and confidence of manually curated disease-gene databases such as SFARI and DDG2P by flagging likely false positive genes. Third, mantis-ml can help clinicians and researchers prioritize which genes to build novel clinical disease-gene cohorts. Often, clinicians and researchers may encounter one or two patients with rare deleterious variants in a gene and submit these genes to tools such as GeneMatcher^40^ to determine if other groups have seen variants linked to similar phenotypes in the same gene. Researchers could focus their efforts on genes with very high mantis-ml percentiles (top 5^th^ percentile, ∼800 genes), re-analyzing existing variants on these genes in addition to reaching out to other groups in a more targeted manner to build disease-gene cohorts. Similarly, mantis-ml could also be used to nominate or de-prioritize genes for deeper functional characterization using model organism or cell-based approaches, as has been done in the Undiagnosed Disease Network’s Model Organism Screening Center^41^. Lastly, we also envision that mantis-ml could be incorporated as gene weights in gene discovery efforts to improve power. The use of genic intolerance to inform gene priors has already led to a greater than 20% increase in ASD gene discovery power ^7^, and mantis-ml’s outperformance of LOEUF across rare variant association studies suggests that it will provide a significant additional boost in power.

The immediate research and clinical impact of these results are significant. First, based on our validation testing, the top 1% of predicted genes from each mantis-ml model provide a high-confidence list of hundreds of likely NDD risk genes for researchers and clinicians across the NDD spectrum. For example, depending on the model, 30-60% of these genes already have publications linking them to phenotypes of interest, a substantial enrichment compared to the rest of the genome. Moreover, the top 1% predicted risk genes are highly enriched compared to the rest of the genome for statistical associations in recent sequencing studies of ASD, DD, and DEE. Second, mantis-ml can help clinicians solve molecular diagnoses. Mantis-ml is a highly accurate NDD risk gene predictor, particularly for genes falling in the top decile of mantis-ml predictions. If a clinician or researcher is presented with a patient with two candidate variants in genes in the top and bottom deciles of mantis-ml risk, depending on the model used, they can have roughly 60-130 times more confidence that the gene in the top decile of risk will be reliably associated with the phenotype of interest. However, we note that the interpretation of the variant effect within any given remains an important challenge in clinical interpretation.

Lastly, while there are several published measures of recessive intolerance ^20–22^, to our knowledge, there are no currently available disease-specific risk predictors for recessive disorders. The discovery of novel recessive disease genes will likely require large sample sizes or access to consanguineous and founder populations given the rarity of homozygous or compound heterozygous pathogenic variants. Until then, mantis-ml’s biallelic models immediately provide a high-confidence assessment of a gene’s probability of being implicated in recessive forms of epilepsy or developmental delay, helping clinicians and researchers solve undiagnosed cases and prioritize genes for deeper functional characterization and gene-matching strategies with other clinicians and patient cohorts. Taken together, our mantis-ml NDD models provide accurate gene risk predictions across the NDD spectrum and illustrate the importance of considering inheritance patterns in generating machine learning-based gene risk predictions.

## Methods

### Seed Gene List Curation

We used SFARI Tier 1 ASD genes (n=207), the highest confidence ranking, as the basis for our monoallelic ASD model. We then reviewed each of the Tier 1 genes to ensure that they were associated with ASD through a monoallelic mechanism and removed genes that had a biallelic mechanism (e.g., *ADSL, ALDH5A1*) or weak evidence of association to ASD based on the most recent large-scale studies of ASD (e.g., *KATNAL2)*. After filtering, we were left with 190 monoallelic ASD seed genes.

For the DD monoallelic and biallelic models, we selected the 832 genes with “Definitive” confidence and “Brain/Cognition” organ involvement from the Developmental Disorder Genotype-Phenotype Database (DD2GP). DD2GP provides mechanism-of-inheritance data for each gene, and we used this information to separate the gene lists into those with monoallelic (N=218) and biallelic inheritance patterns (N=449). For the DD monoallelic model, we combined the DD2GP monoallelic genes with 199 genome-wide significant genes from the largest trio exome sequencing study of DD^2^, resulting in a total of 417 monoallelic seed genes for DD.

For DEE, we first selected genes from OMIM^33^ with the specific phenotype of “Developmental and Epileptic Encephalopathy” and stratified them based on their pattern of inheritance. We then combined these genes with the list of clinically curated DEE genes from the most recent Epi25k study of epilepsy,^1^ stratified by inheritance. Lastly, we added additional genes from OMIM that had robust evidence for causing epileptic encephalopathy by conducting an advanced search for “epileptic encephalopathy” and manually curating the strength of the associated literature. Genes that were associated with epilepsy but not robustly associated with epileptic encephalopathy were not included as seed genes (e.g., *DEPDC5)* as we aimed to train the model on the most severe forms of epilepsy.

### Fetal cortex scRNA-seq analysis

We downloaded single-cell RNA expression data generated from human fetal cortical samples as described in a prior publication^26^. The data, which include four samples from an 8-week span during mid-gestation, consisted of 57,868 single-cell transcriptomes. Using the same cell identity annotations from the original publication, we calculated the module score for all cells using Seurat’s AddModuleScore function with the seed gene list for each NDD as the input feature ^27,42^. We then Z-score normalized these module scores to evaluate the relative expression of disease-associated genes between cell clusters. We also calculated the average unique molecular identifier (UMI) counts of all genes per cell type per age of tissue. These were used as features in the machine learning models.

### Fetal cortex scRNA-sequencing models

To demonstrate the baseline power of fetal cortex single-cell RNA-sequencing data as a predictor of NDD risk genes, we evaluated the performance of a Random Forest model for each set of curated seed genes. Due to the intrinsic imbalance between the positively labeled and unlabeled genes, machine learning models can quickly become biased towards the majority class. Reducing the size of the overrepresented class can help diminish the likelihood of a model overfitting, providing more accurate predictions. We thus created balanced datasets for each inheritance-specific phenotype seed gene list consisting of all positively labeled genes and a random subset of unlabeled genes, with the resulting dataset containing a ratio of positively labeled to unlabeled genes of 1:1.5. Next, we performed zero imputation and removed highly correlated features (Pearson’s *r* > 0.95). We used the scikit-learn library in Python to construct the Random Forest Model with the default parameters. Using 5-fold cross-validation, we evaluated the performance of the random forest model for each dataset by calculating the area under the receiver operater curve. Additionally, we compared the performance of the scRNA-seq expression models to models trained on intolerance metrics, including missense Z, RVIS, LOEUF, and pREC ^10,13,20^.

The Boruta algorithm is an iterative feature selection method, using the Random Forest algorithm during learning, that determines if a feature has a statistically robust predictive power. Unlike other feature selection methods where features are compared against each other, Boruta compares each feature against randomized versions of the original feature set called “shadow” features. Features achieving less significant importance than the “shadow” features are progressively eliminated. Eventually, a “confirmed” set of features (i.e., features that are considered predictive) are identified and ranked based on Z-scores representing importance scores. We employed the Boruta algorithm in R to evaluate the feature importance of our fetal cortex scRNA-seq data. We repeated this step for each model using the default parameters and the corresponding balanced dataset with all fetal cortex scRNA-seq features.

### Mantis-ml

The mantis-ml framework has been previously described in detail ^25^. Briefly, mantis-ml is a semi-supervised machine learning framework for the prediction of possible novel disease-associated genes. Following its initial setup, disease/phenotype terms of interest provided to mantis-ml are used to automatically extract associated known disease genes and relevant features. In this manuscript, given the relative importance of using high confidence seed genes, we elected to manually curate our seed genes. Next, mantis-ml annotates all genes in the genome with several hundred diverse features. The semi-supervised learning method employed by mantis-ml infers the risk of a gene’s association with a specific phenotype based on the similarity between the feature signature of the gene and that of the seed genes, as captured by hundreds or thousands of balanced sets comprising of known and unlabeled genes. Mantis-ml then generates exome-wide gene-level risk prediction probabilities and their corresponding percentiles for the phenotype of interest.

We instituted several key improvements to the original mantis-ml framework to improve performance for NDD risk gene prediction. First, we integrated fetal human cortex scRNA-seq data containing the average gene expression for each major cell type over four different developmental stages. We expanded on the previous collection of gene intolerance metrics in mantis-ml by adding the gene variation intolerance rank (GeVIR)^29^, which performs well for smaller genes and missense intolerant genes. We also added GeVIR’s LOEUF-joined derivative, ViRLoF, which has been shown to outperform LOEUF alone in prioritizing NDD risk genes. Lastly, we included GeVIR’s fold-enrichment scores for autosomal dominant and recessive modes of inheritance for each gene. Gene Ontology (GO) terms are a powerful tool for describing the relationship between a gene/gene product and its functional, molecular, and spatial properties. The original mantis-ml framework incorporated GO terms by applying a pattern search using the disease/phenotype input terms and collapsing the number of associations between a gene and the matched GO terms into a new, one-hot encoded feature per input term. We now expand the GO feature set by also including the top 20 individual GO terms that seed genes are most enriched for compared to the rest of the exome (quantified via Fisher’s exact test), further increasing the strength of the mantis-ml feature set.

We used fixed configuration and classifier parameters for each input seed gene list and their corresponding disease/phenotype terms of interest, as described in the original mantis-ml publication (**Table S24**). Prior to model training and inference, mantis-ml automatically performs preprocessing and exploratory data analysis (EDA). The initial preprocessing step of mantis-ml performs feature filtering by calculating Pearson’s correlation coefficient between all features and dropping those with correlations above a defined threshold. To prevent the removal of valuable scRNA-seq features due to genes with low or non-existent expression in fetal cortex tissue, we specified a high correlation threshold of 0.95 (default = 0.8). We retained the default mantis-ml parameters for the remainder of the preprocessing and EDA steps.

For the stochastic semi-supervised component of mantis-ml, we utilized the random forest and extreme gradient boosting (XGBoost) classifiers due to their superior performance in the original paper. First, mantis-ml generated balanced datasets (*M*) containing a ratio of randomly selected positively labeled genes to randomly selected unlabeled genes equal to 1:1.5, with the positively labeled genes containing only 80% of known disease-associated genes. For each balanced dataset, mantis-ml performed stratified *k*-fold cross-validation with out-of-bag prediction using k = 10 folds. After prediction probabilities are generated for the entire gene space, mantis-ml repeats this process a total of 10 times (L) by creating new balanced datasets and performing stratified k-fold cross-validation. Finally, mantis-ml generated a ranked candidate gene list by computing the mean prediction probability and corresponding percentile score for each gene.

### Validation of mantis-ml using rare variant association study summary statistics

We tested whether top predicted monoallelic mantis-ml risk genes were enriched for genes with statistical support from recent large-scale sequencing studies. We obtained summary statistics from the largest and most recently available studies of ASD and DD^6^ and epileptic encephalopathy ^1^. All three of these association studies only included dominant models. Thus, we tested for enrichment across the three relevant monoallelic mantis-ml models. Using a two-tailed Fisher’s exact test, we calculated the enrichment of the top 5^th^ percentile of mantis-ml predictions among nominally significant genes (p<0.01) from each of the three association studies. Fu et al. did not include the X-chromosome in their test, so excluded X-chromosome genes in the enrichment tests for ASD and DD. We also calculated the enrichment of genes highly intolerant to loss-of-function variation (LOEF top 5^th^ percentile). Due to potential concerns of circularity, we repeated these enrichment tests excluding seed genes.

### Validation of mantis-ml with clinically curated gene lists

We downloaded clinically curated gene lists from Simons Foundation Autism Research Initiative (SFARI) for ASD (download date: 01/18/2022) and DECIPHER’s Developmental Disorder Genotype – Phenotype Database (DDG2P) for DD (download date: 12/16/2021). SFARI currently provides three Tiers of confidence and DDG2P provides five including Definitive (our seed genes), Strong, Moderate, Limited, and Relevant Disease and Incidental Finding (RD/IF). For our analysis, we only used Strong and Limited as there were too few genes with Moderate and RD/IF classifications. We then plotted the distribution of monoallelic ASD mantis-ml percentiles for Tier 1 (seed genes), Tier 2, and Tier 3 ASD genes and compared them to the distribution to the rest of the genes in the exome not included in Tiers 1, 2, and 3. We repeated the same procedures for the DDG2P “Definitive” (seed genes), “Strong”, and “Limited” evidence genes, stratified by monoallelic and biallelic inheritance with the distribution of their respective mantis-ml risk percentiles.

We then evaluated the degree of enrichment of genes in the top 5^th^ percentile of mantis-ml predictions across each tier / category using a two-tailed Fisher’s exact test. We also calculated the enrichment for two intolerance metrics: LOEUF and pREC (for biallelic/recessive lists). For each enrichment test, we compared genes within each tier/category to genes in the rest of the protein-coding genome that were not contained in any other category.

### Validation of mantis-ml using an automated literature search with AMELIE

Further validation of mantis-ml results was performed using AMELIE (Automatic Mendelian Literature Evaluation)^32^. Briefly, AMELIE uses natural language processing to identify manuscripts from the extant literature with a phenotype match for genes of interest. For each manuscript with a gene-phenotype match, AMELIE reports a phenotypic match score based on the strength of the match of the language in the manuscript with the Human Phenotype Ontology input term. A match of 100% represents a perfect match for a gene and given phenotype and lower phenotypic match percentiles are given for related descendant phenotypes in the Human Phenotype Ontology (HPO)^43^.

For each model, we generated genome-wide AMELIE phenotype match scores in a two-step process. Using the default parameters, we ran AMELIE with the HPO terms HP:0000729 (Autistic Behavior), HP:0001250 (Seizures), and HP:0012759 (Neurodevelopmental abnormality) for ASD, DEE, and DD, respectively. We repeated this process with the inheritance mode parameter set to “dominant”. Although AMELIE does not permit the use of “recessive” as an inheritance mode filter, it assigns both recessive and dominant scores based on the context of an article. The dominant inheritance mode instructs AMELIE to avoid returning articles for genes with higher recessive scores. Therefore, we treated the non-union of genes between the non-specified inheritance and dominant runs as our recessive set of AMELIE scores (**Tables S25-29**). For each set of mantis-ml ranked predictions, we annotated genes with their corresponding phenotypic match score and removed the seed genes from the dataset. We then used Fisher’s exact test to determine the enrichment of at least one publication with a 100% phenotypic match score in each mantis-ml decile across all models. We repeated this process using the most stringent level of evidence that AMELIE allows (five or more publications with 100% phenotypic match scores) to evaluate mantis-ml’s performance with the highest confidence gene-phenotype matches.

## Acknowledgments

We thank Dr. Huda Zoghbi for useful discussions and valuable feedback. This work was supported by grant NIH NINDS F32 NS127854 (R.S.D.) and K23MH121669 (A.W.Z.).

## Data Availability

The mantis-ml predictions for all five NDD models are available as supplementary files and through a publicly available browser: https://nddgenes.com.

## Code Availability

The mantis-ml code is available on GitHub (https://github.com/astrazeneca-cgr-publications/mantis-ml-release).

## References

1. Feng, Y.-C. A. et al. Ultra-Rare Genetic Variation in the Epilepsies: A Whole-Exome Sequencing Study of 17,606 Individuals. The American Journal of Human Genetics 105, 267–282 (2019).

2. Kaplanis, J. et al. Evidence for 28 genetic disorders discovered by combining healthcare and research data. Nature 586, 757–762 (2020).

3. McRae, J. F. et al. Prevalence and architecture of de novo mutations in developmental disorders. Nature 542, 433–438 (2017).

4. Motelow, J. E. et al. Sub-genic intolerance, ClinVar, and the epilepsies: A whole-exome sequencing study of 29,165 individuals. The American Journal of Human Genetics 108, 965–982 (2021).

5. Allen, A. S. et al. De novo mutations in epileptic encephalopathies. Nature 501, 217–221 (2013).

6. Fu, J. M. et al. Rare coding variation illuminates the allelic architecture, risk genes, cellular expression patterns, and phenotypic context of autism. http://medrxiv.org/lookup/doi/10.1101/2021.12.20.21267194 (2021) doi:10.1101/2021.12.20.21267194.

7. Satterstrom, F. K. et al. Large-Scale Exome Sequencing Study Implicates Both Developmental and Functional Changes in the Neurobiology of Autism. Cell 180, 568-584.e23 (2020).

8. Srivastava, S. et al. Meta-analysis and multidisciplinary consensus statement: exome sequencing is a first-tier clinical diagnostic test for individuals with neurodevelopmental disorders. Genet Med 21, 2413–2421 (2019).

9. Iossifov, I. et al. The contribution of de novo coding mutations to autism spectrum disorder. Nature 515, 216–221 (2014).

10. Petrovski, S., Wang, Q., Heinzen, E. L., Allen, A. S. & Goldstein, D. B. Genic Intolerance to Functional Variation and the Interpretation of Personal Genomes. PLoS Genet 9, e1003709 (2013).

11. Dhindsa, R. S., Copeland, B. R., Mustoe, A. M. & Goldstein, D. B. Natural Selection Shapes Codon Usage in the Human Genome. The American Journal of Human Genetics 107, 83– 95 (2020).

12. Traynelis, J. et al. Optimizing genomic medicine in epilepsy through a gene-customized approach to missense variant interpretation. Genome Res 27, 1715–1729 (2017).

13. Samocha, K. E. et al. A framework for the interpretation of de novo mutation in human disease. Nat Genet 46, 944–950 (2014).

14. Vitsios, D., Dhindsa, R. S., Middleton, L., Gussow, A. B. & Petrovski, S. Prioritizing non-coding regions based on human genomic constraint and sequence context with deep learning. Nat Commun 12, 1504 (2021).

15. Palmer, D. S. et al. Exome sequencing in bipolar disorder identifies AKAP11 as a risk gene shared with schizophrenia. Nat Genet 54, 541–547 (2022).

16. Singh, T. et al. Rare coding variants in ten genes confer substantial risk for schizophrenia. Nature 604, 509–516 (2022).

17. Zoghbi, A. W. et al. High-impact rare genetic variants in severe schizophrenia. Proc Natl Acad Sci USA 118, e2112560118 (2021).

18. Halvorsen, M. et al. Exome sequencing in obsessive–compulsive disorder reveals a burden of rare damaging coding variants. Nat Neurosci 24, 1071–1076 (2021).

19. iPSYCH-Broad Consortium et al. Autism spectrum disorder and attention deficit hyperactivity disorder have a similar burden of rare protein-truncating variants. Nat Neurosci 22, 1961–1965 (2019).

20. Karczewski, K. J. et al. The mutational constraint spectrum quantified from variation in 141,456 humans. Nature 581, 434–443 (2020).

21. Balick, D. J., Jordan, D. M., Sunyaev, S. & Do, R. Overcoming constraints on the detection of recessive selection in human genes from population frequency data. The American Journal of Human Genetics 109, 33–49 (2022).

22. Hsu, J. S. et al. Inheritance-mode specific pathogenicity prioritization (ISPP) for human protein coding genes. Bioinformatics 32, 3065–3071 (2016).

23. Krishnan, A. et al. Genome-wide prediction and functional characterization of the genetic basis of autism spectrum disorder. Nat Neurosci 19, 1454–1462 (2016).

24. Liu, L. et al. DAWN: a framework to identify autism genes and subnetworks using gene expression and genetics. Molecular Autism 5, 22 (2014).

25. Vitsios, D. & Petrovski, S. Mantis-ml: Disease-Agnostic Gene Prioritization from High-Throughput Genomic Screens by Stochastic Semi-supervised Learning. The American Journal of Human Genetics 106, 659–678 (2020).

26. Trevino, A. E. et al. Chromatin and gene-regulatory dynamics of the developing human cerebral cortex at single-cell resolution. Cell 184, 5053-5069.e23 (2021).

27. Tirosh, I. et al. Dissecting the multicellular ecosystem of metastatic melanoma by single-cell RNA-seq. Science 352, 189–196 (2016).

28. Polioudakis, D. et al. A Single-Cell Transcriptomic Atlas of Human Neocortical Development during Mid-gestation. Neuron 103, 785-801.e8 (2019).

29. Abramovs, N., Brass, A. & Tassabehji, M. GeVIR is a continuous gene-level metric that uses variant distribution patterns to prioritize disease candidate genes. Nat Genet 52, 35–39 (2020).

30. Abrahams, B. S. et al. SFARI Gene 2.0: a community-driven knowledgebase for the autism spectrum disorders (ASDs). Molecular Autism 4, 36 (2013).

31. Bragin, E. et al. DECIPHER: database for the interpretation of phenotype-linked plausibly pathogenic sequence and copy-number variation. Nucl. Acids Res. 42, D993–D1000 (2014).

32. Birgmeier, J. et al. AMELIE speeds Mendelian diagnosis by matching patient phenotype and genotype to primary literature. Sci. Transl. Med. 12, eaau9113 (2020).

33. Hamosh, A., Scott, A. F., Amberger, J. S., Bocchini, C. A. & McKusick, V. A. Online Mendelian Inheritance in Man (OMIM), a knowledgebase of human genes and genetic disorders. Nucleic acids research 33, D514–D517 (2005).

34. Bhalla, K. et al. Alterations in CDH15 and KIRREL3 in patients with mild to severe intellectual disability. Am J Hum Genet 83, 703–713 (2008).

35. Kaname, T. et al. Mutations in CD96, a member of the immunoglobulin superfamily, cause a form of the C (Opitz trigonocephaly) syndrome. Am J Hum Genet 81, 835–841 (2007).

36. Darlow, J. M., McKay, L., Dobson, M. G., Barton, D. E. & Winship, I. On the origins of renal cell carcinoma, vesicoureteric reflux and C (Opitz trigonocephaly) syndrome: A complex puzzle revealed by the sequencing of an inherited t (2; 3) translocation. Eur. J. Hum. Genet 21, 145 (2013).

37. Urreizti, R. et al. Screening of CD96 and ASXL1 in 11 patients with Opitz C or Bohring– Opitz syndromes. American Journal of Medical Genetics Part A 170, 24–31 (2016).

38. Chan, C. J. et al. The receptors CD96 and CD226 oppose each other in the regulation of natural killer cell functions. Nature immunology 15, 431–438 (2014).

39. Schaaf, C. P. et al. A framework for an evidence-based gene list relevant to autism spectrum disorder. Nat Rev Genet 21, 367–376 (2020).

40. Sobreira, N., Schiettecatte, F., Valle, D. & Hamosh, A. GeneMatcher: A Matching Tool for Connecting Investigators with an Interest in the Same Gene. Human Mutation 36, 928–930 (2015).

41. Undiagnosed Diseases Network et al. Model organisms contribute to diagnosis and discovery in the undiagnosed diseases network: current state and a future vision. Orphanet J Rare Dis 16, 206 (2021).

42. Stuart, T. et al. Comprehensive Integration of Single-Cell Data. Cell 177, 1888-1902.e21 (2019).

43. Robinson, P. N. et al. The Human Phenotype Ontology: A Tool for Annotating and Analyzing Human Hereditary Disease. The American Journal of Human Genetics 83, 610–615 (2008).

