## Supplementary Information for "Genome-wide prediction of dominant and recessive neurodevelopmental disorder risk genes"

**The PDF file includes:**

Figure. S1. scRNA-sequencing model Boruta feature importance.

Figure. S2. ASD Mantis-ml Boruta feature importance

Figure. S3. DD Mantis-ml Boruta feature importance

Figure. S4. DEE Mantis-ml Boruta feature importance

Figure S5. Comparison of inheritance-informed and non-inheritance informed mantis-ml model performance across rare variant association studies.

Figure S6. Comparison of inheritance-informed and non-inheritance informed mantis-ml model performance across rare variant association studies and clinically curated gene list.

Figure S7. Frequency of clinically curated genes supported by one or more publications in the top and bottom mantis-ml prediction percentiles.

Figure S8. Distribution of statistical support for clinically curated genes in mantis-ml in the top and bottom mantis-ml prediction percentiles

Figure S9. Distribution of genes with one or more phenotype-associated publications in mantis-ml prediction deciles.

Figure S10. Distribution of genes with five or more phenotype-associated publications in mantis-ml prediction deciles.

Figure S11. Comparison of inheritance-informed and non-inheritance informed model enrichment of genes with 100% phenotype match in published literature stratified by mantis-ml decile.

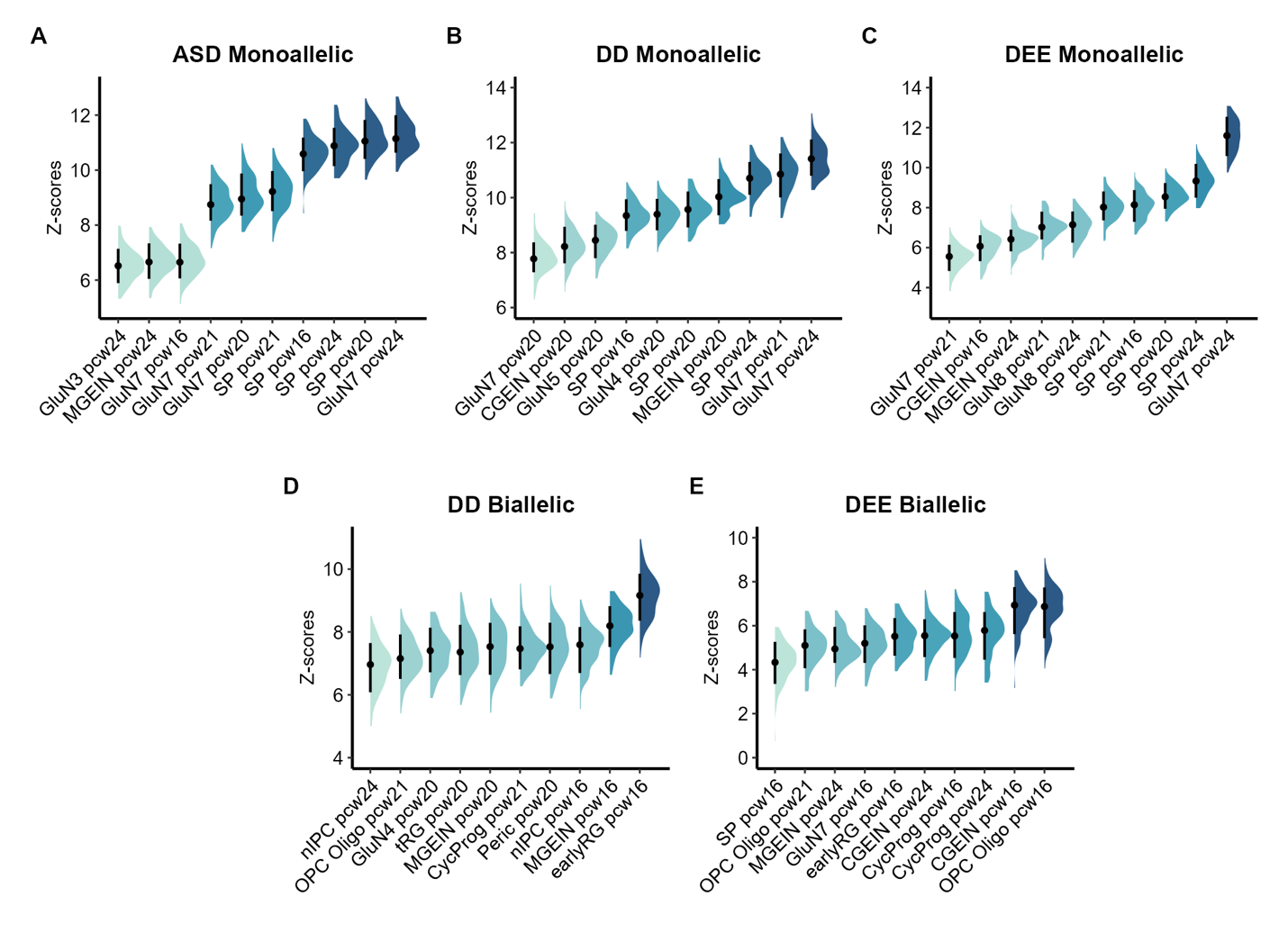
**Figure S1. Feature importances of scRNA-seq-derived random forest NDD models.** Top 10 feature importances derived from the Boruta algorithm for five random forest models trained on human fetal scRNA-seq data for monoallelic ASD **(A)**, monoallelic DD **(B)**, monoallelic DEE **(C)**, biallelic DD **(D)**, and biallelic DEE **(E).**

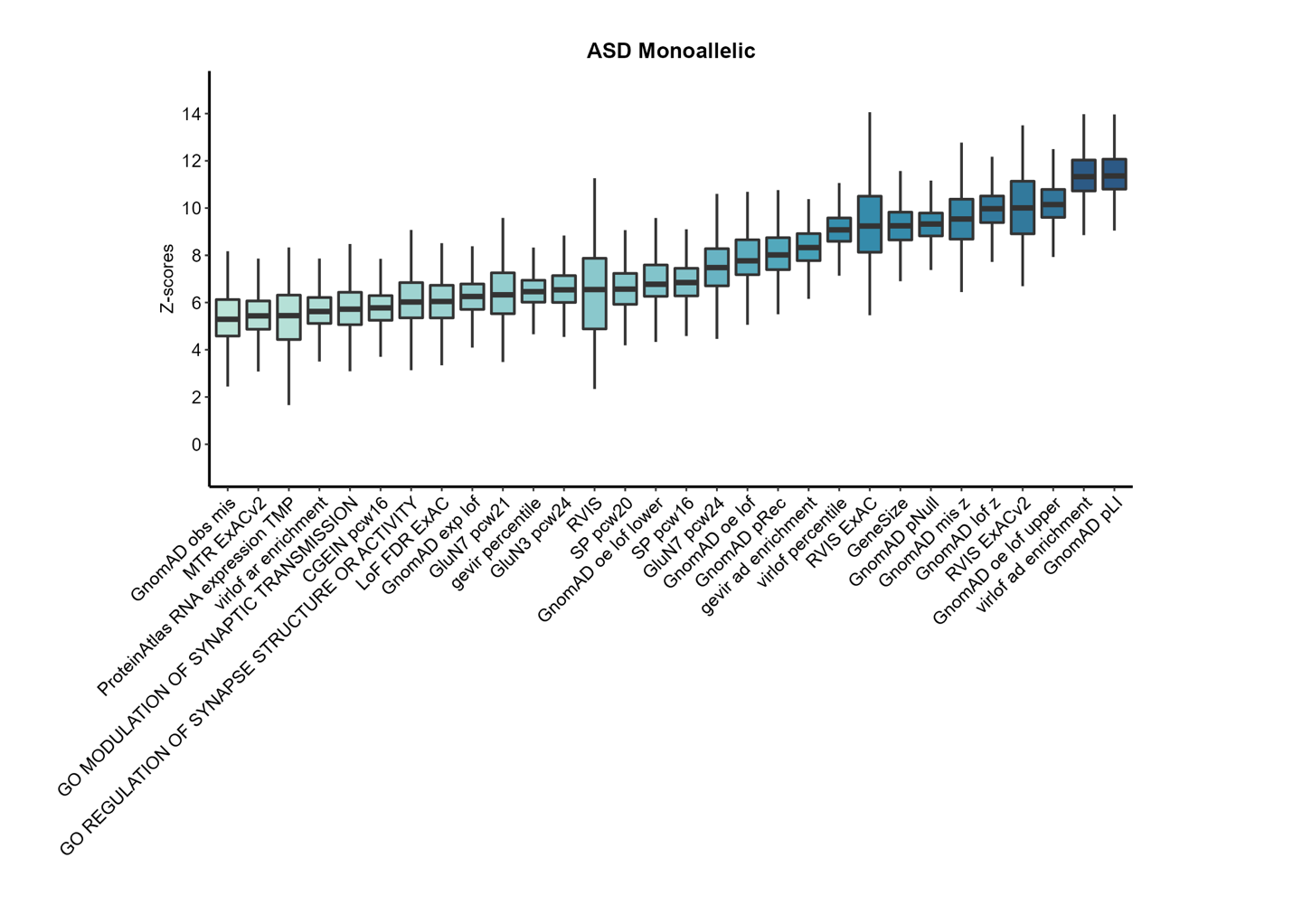

**Figure S2. Monoallelic ASD mantis-ml feature importance.** Top 30 monoallelic ASD mantis-ml feature importances based on Boruta.

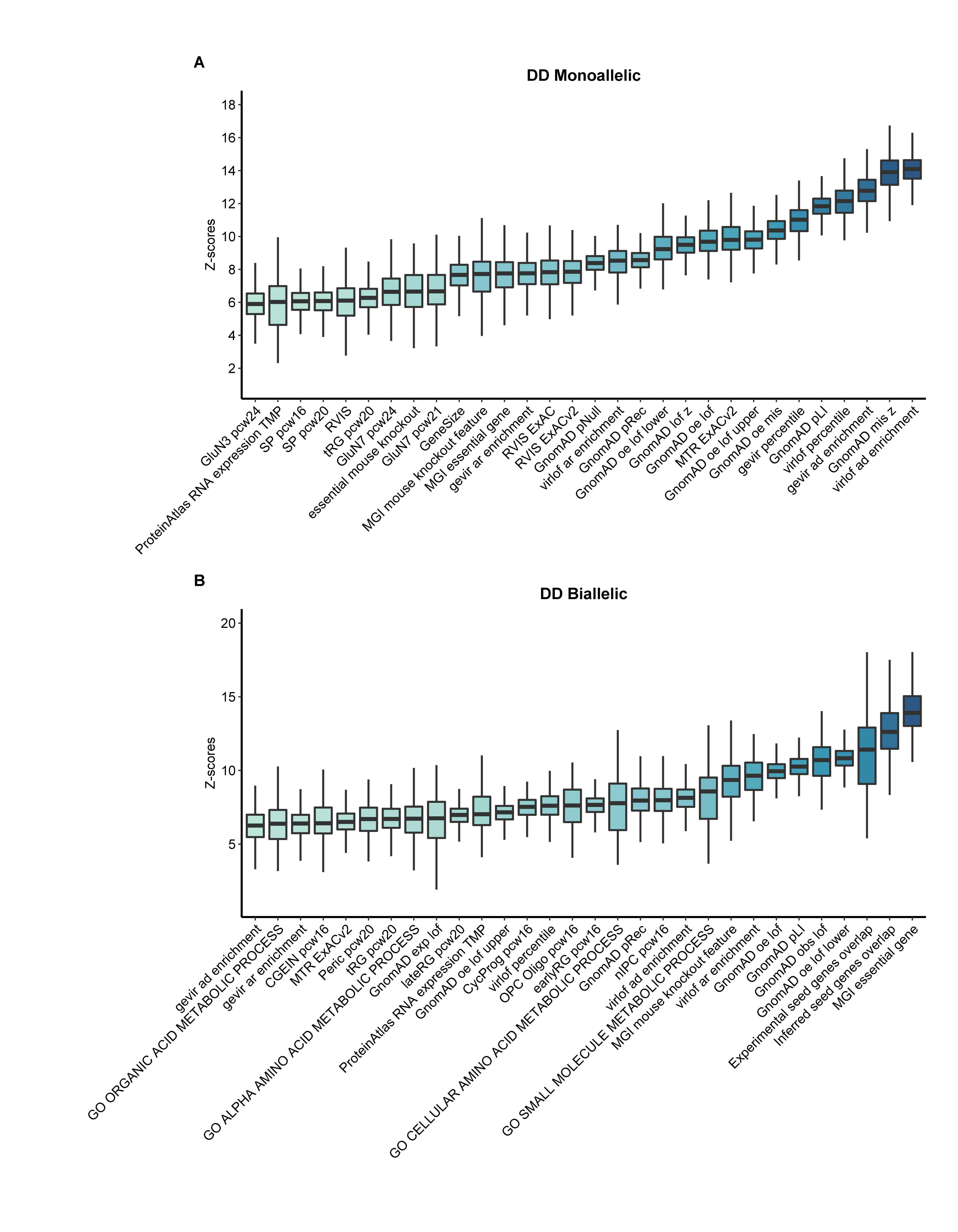

**Figure S3. Mantis-ml feature importances for DD.** Top 30 monoallelic mantis-ml feature importances based on Boruta for **(A)** monoallelic DD and **(B)** biallelic DD**.**

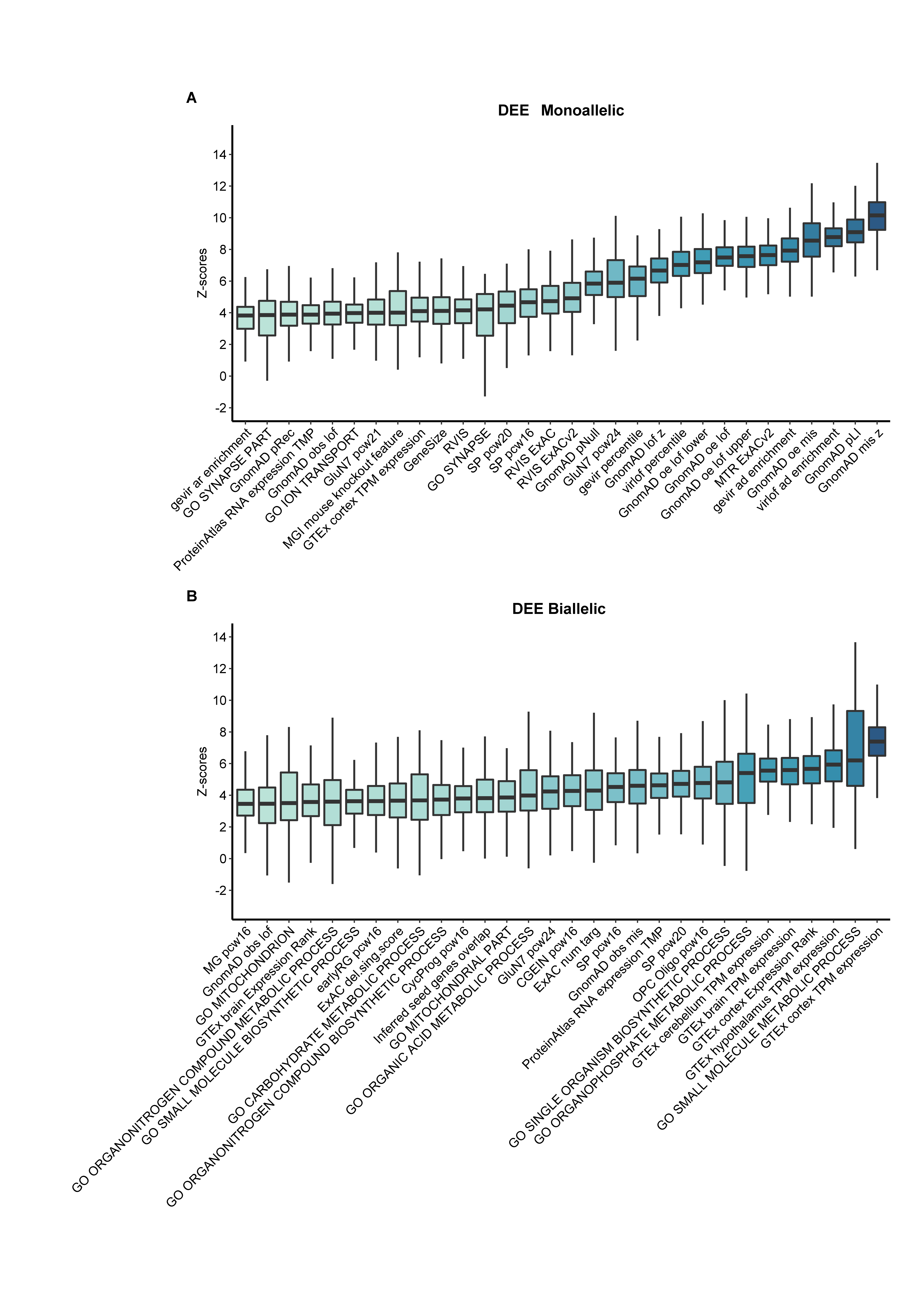

**Figure S4. Mantis-ml feature importances for DEE.** Top 30 monoallelic mantis-ml feature importances based on Boruta for **(A)** monoallelic DEE and **(B)** biallelic DEE**.**

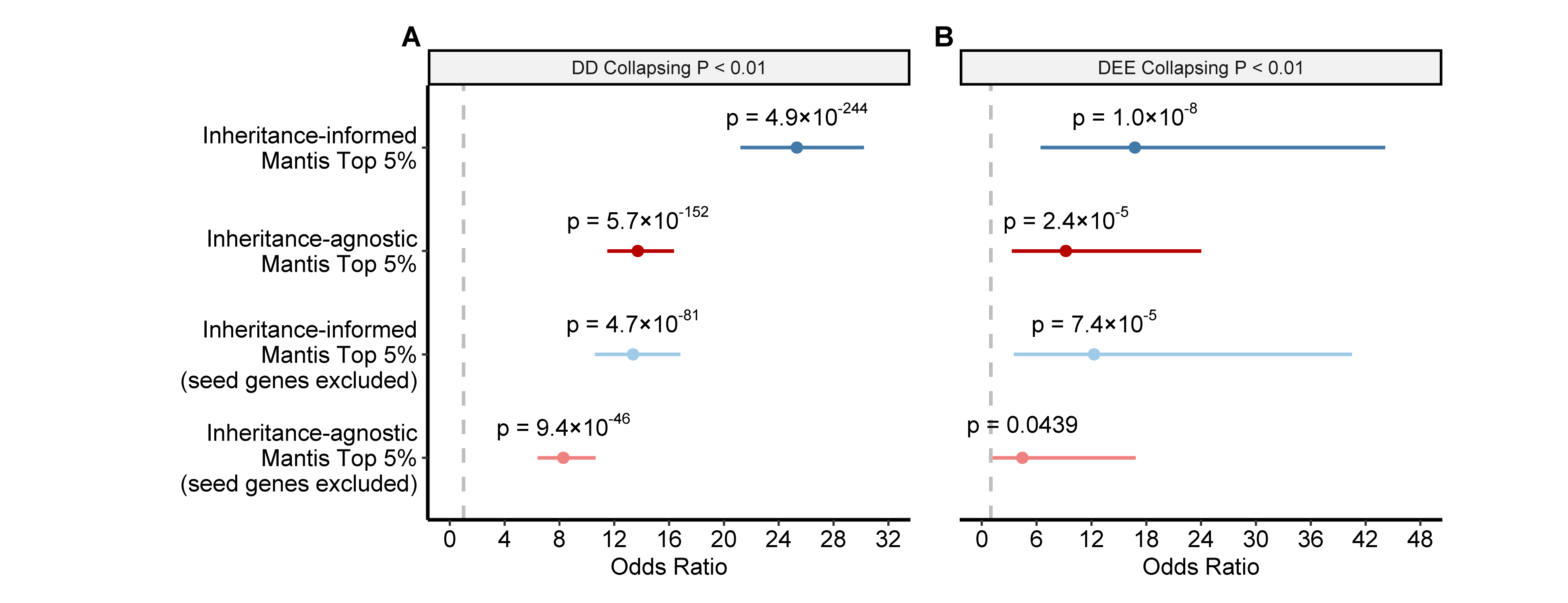

**Figure S5. Comparison of inheritance-informed and inheritance-agnostic mantis-ml model performance across rare variant association studies. (A)** Enrichment of nominally significant (p<0.01) genes from a DD gene-based collapsing analysis among the top 5^th^ percentile of predictions from mantis-ml models trained on only monoallelic DD seed genes (“inheritance-informed”) versus all DD seed genes (“inheritance-agnostic”). **(B)** Enrichment of nominally significant genes from DEE gene-based collapsing analysis among the top 5^th^ percentile of mantis-ml models based on either inheritance-informed or inheritance-agnostic DEE seed genes.

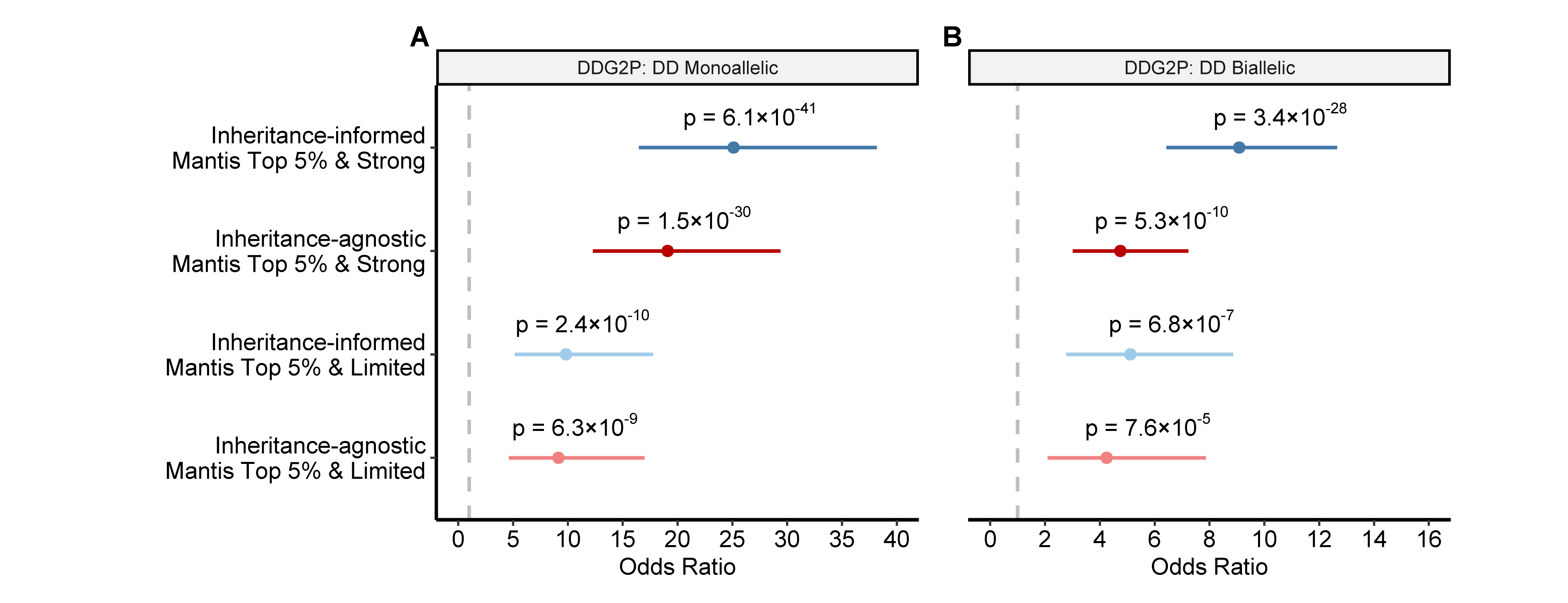

**Figure S6. Comparison of inheritance-informed and inheritance-agnostic mantis-ml model performance across clinically curated DD gene lists. (A)** Enrichment of top 5^th^ percentile predictions from the monoallelic DD mantis-ml model versus an inheritance-agnostic DD mantis-ml model among DDG2P “Strong” and “Limited” monoallelic gene lists. **(B)** Enrichment of top 5^th^ percentile predictions from the biallelic DD mantis-ml model versus the inheritance-agnostic DD mantis-ml model among DDG2P “Strong” and “Limited” biallelic gene lists.

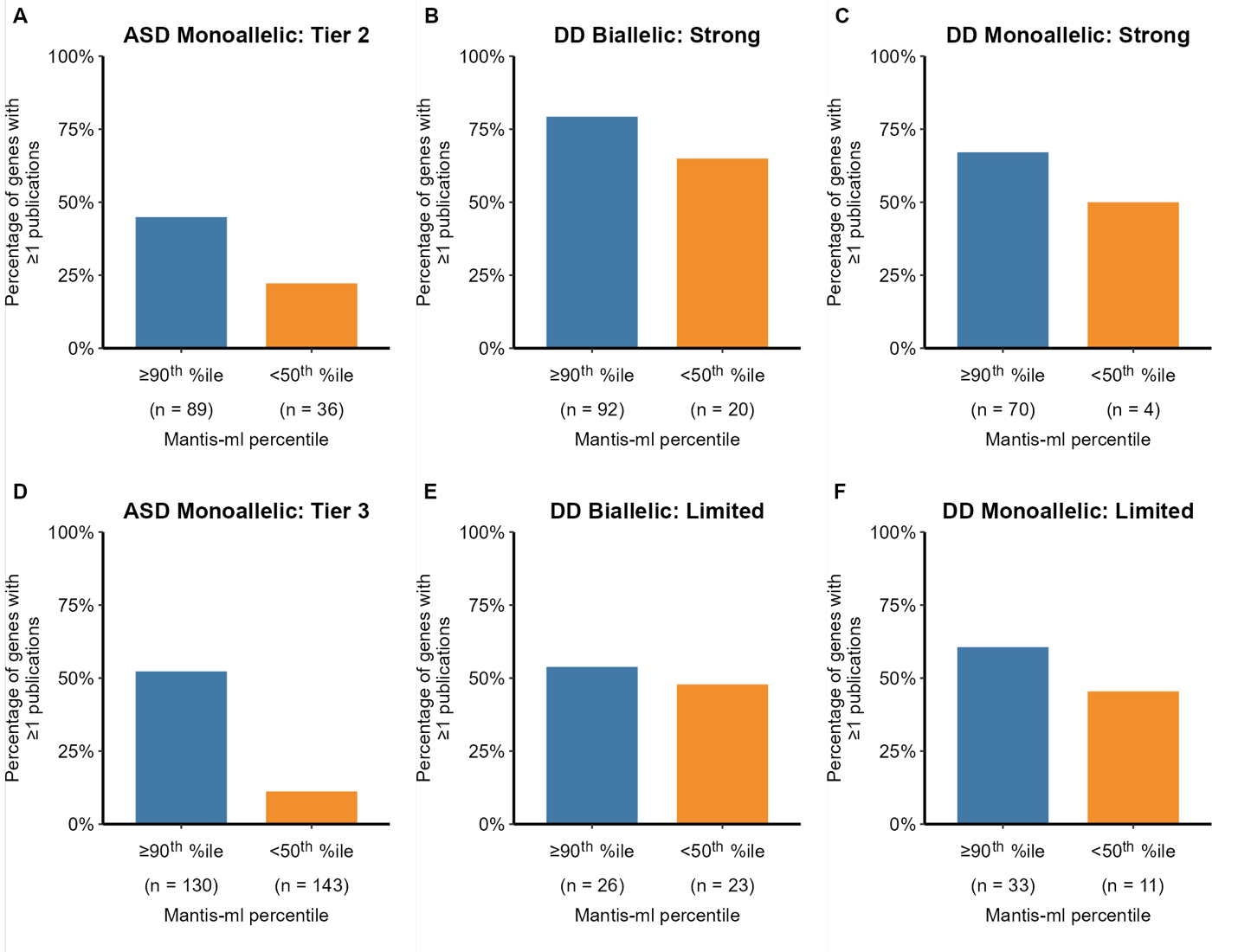

**Figure S7. Percentage of SFARI and DDG2P genes with supporting publications detected via AMELIE, stratified by mantis-ml probabilities**. **(A)** Percentage of Tier 2 SFARI ASD genes

with at least one supporting publication detected via AMELIE falling into the top 10^th^ percentile of monoallelic ASD mantis-ml predictions versus the bottom 50^th^ percentile. **(B)** Percentage of DDG2P “Strong” biallelic DD genes with at least one supporting publication stratified by the mantis-ml biallelic DD percentiles. **(C)** Percentage of DDG2P “Strong” monoallelic DD genes with at least one supporting publication stratified by the mantis-ml monoallelic DD percentiles. **(D)** Percentage of SFARI Tier 3 monoallelic ASD genes with at least one supporting publication stratified by the mantis-ml monoallelic ASD percentiles. **(E)** Percentage of DDG2P “Limited” biallelic DD genes with at least one supporting publication stratified by the mantis-ml biallelic DD percentiles. **(F)** Percentage of DDG2P “Limited” monoallelic DD genes with at least one supporting publication stratified by the mantis-ml monoallelic DD percentiles.

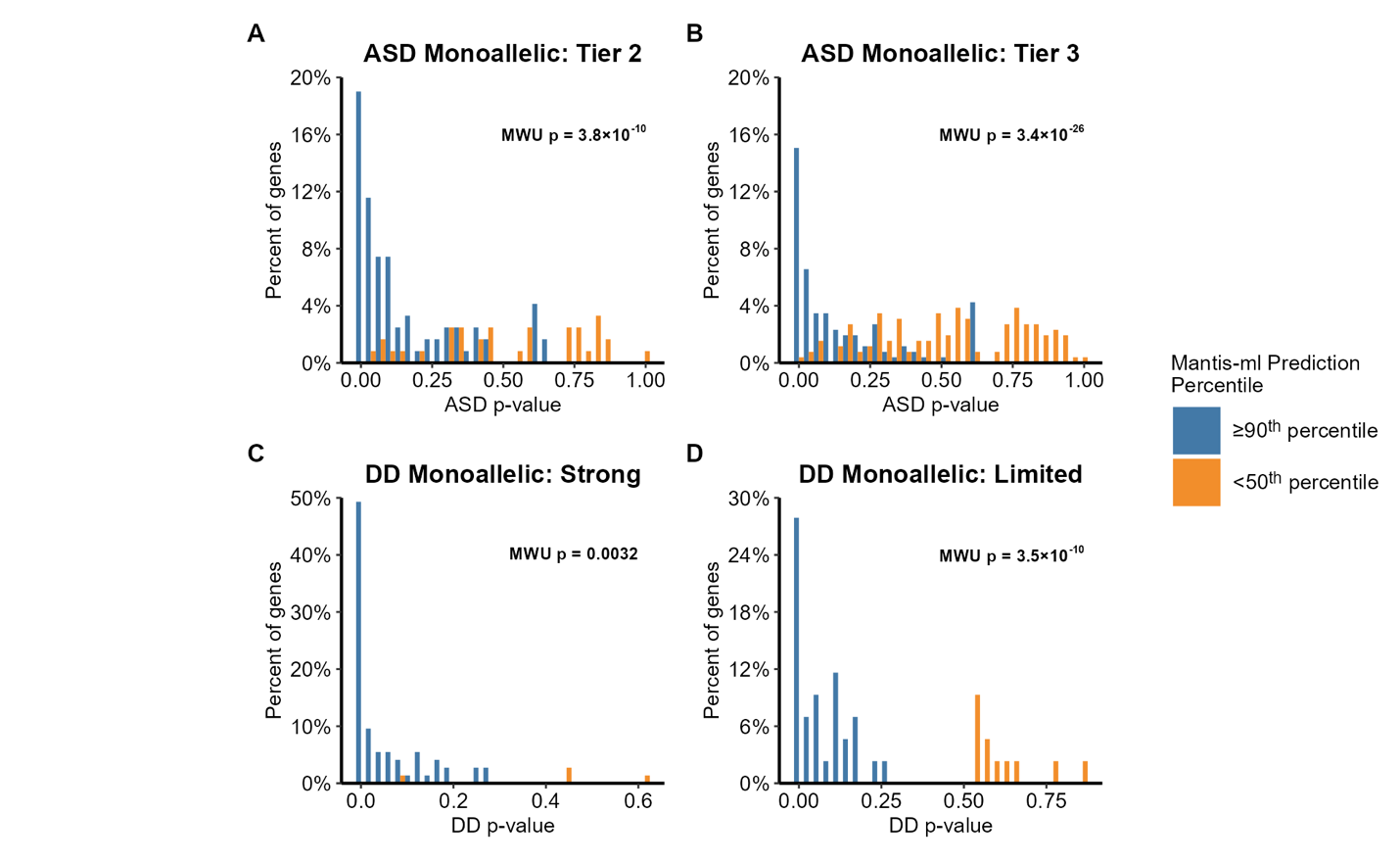

**Figure S8. Distribution of p-values from rare variant association studies for top and bottom mantis-ml predictions.** (**A**) P-value distributions of genes in SFARI Tier 2 based on a recent ASD collapsing analysis, colored by top (>=90^th^ percentile) and bottom (<50^th^ percentile) mantis-ml monoallelic ASD predictions. **(B)** Same as (A) for Tier 3 SFARI genes. **(C)** P-value distributions of genes in the DDG2P “Strong” monoallelic gene list based on a recent DDD collapsing analysis, colored by top (>=90^th^ percentile) and bottom (<50^th^ percentile) mantis-ml monoallelic DD predictions. **(D)** Same as (C) for DDG2P “Limited” monoallelic genes.

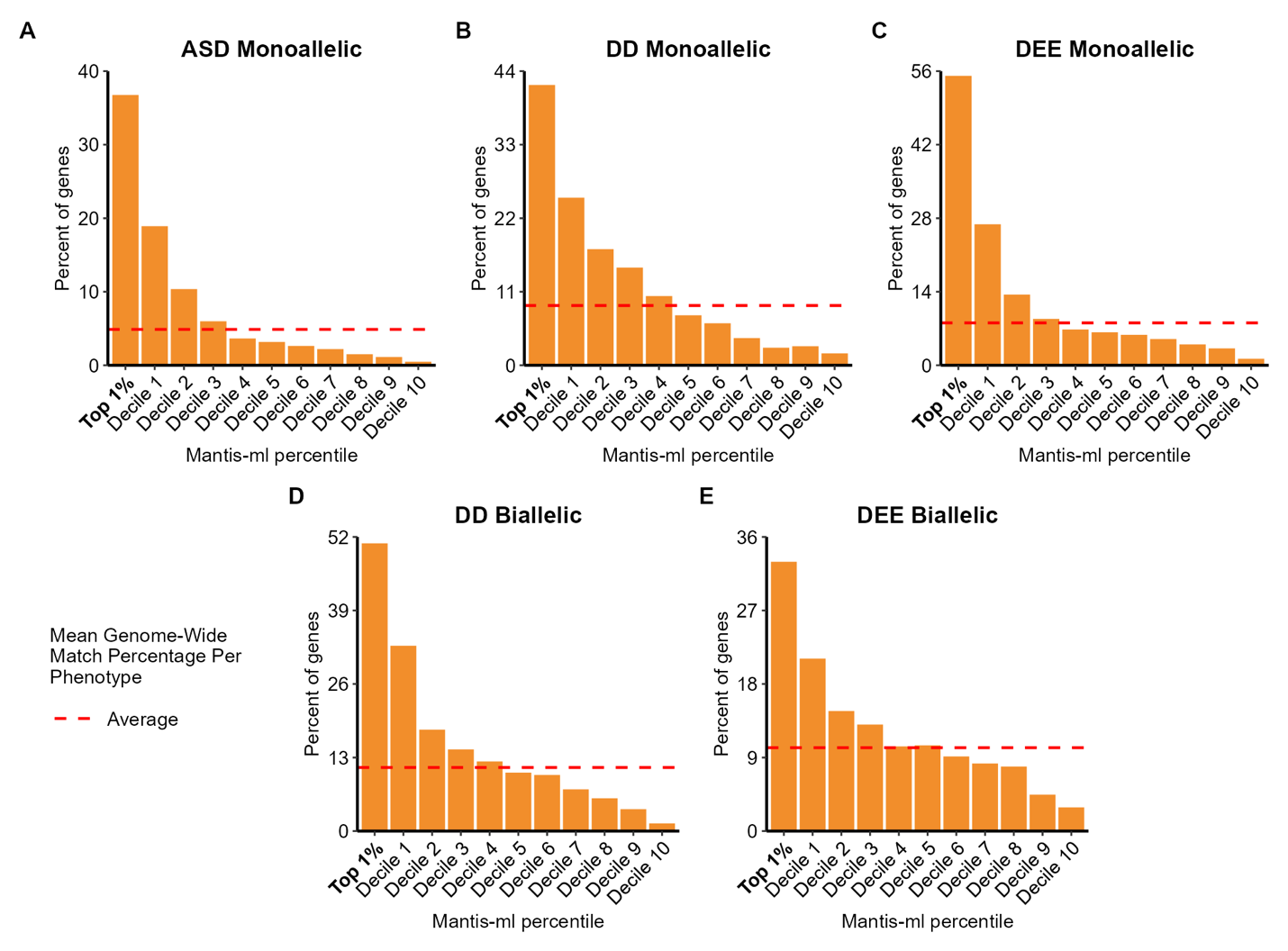

**Figure S9. Distribution of genes with one or more phenotype-associated publications in neurodevelopmental disorder mantis-ml prediction deciles.** (**A-C**) Percentage of genes per mantis-ml decile with at least one phenotype-associated publication detected via AMELIE for monoallelic (A) ASD, (B) DD, and (C) DEE. **(D, E)** Percentage of genes per mantis-ml decile with at least one phenotype-associated publication detected via AMELIE for biallelic (D) DD and (E) DEE.

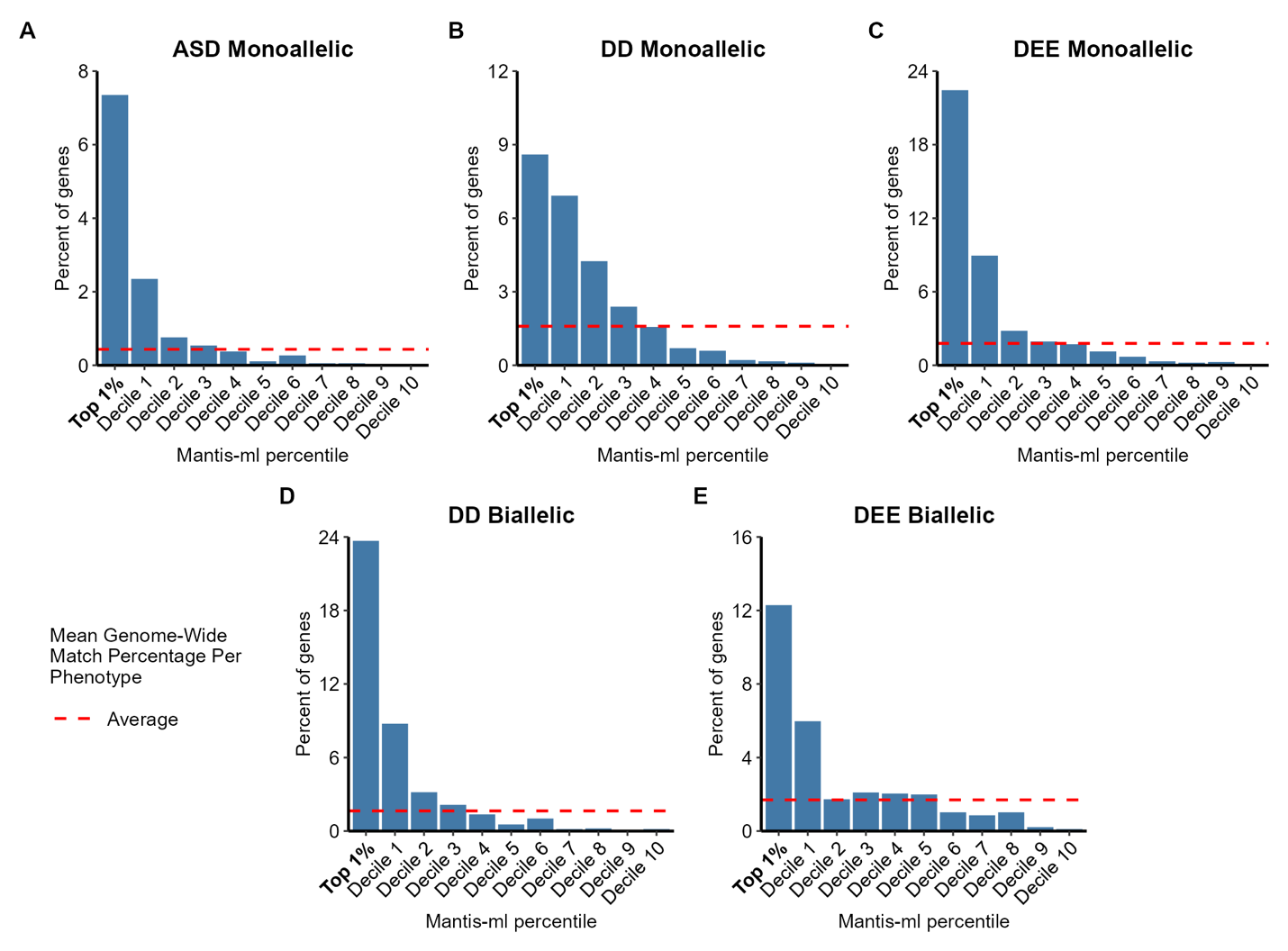

**Figure S10. Distribution of genes with five or more phenotype-associated publications in neurodevelopmental disorder mantis-ml prediction deciles.** (**A-C**) Percentage of genes per mantis-ml decile with at least five phenotype-associated publications detected via AMELIE for monoallelic (A) ASD, (B) DD, and (C) DEE. **(D, E)** Percentage of genes per mantis-ml decile with at least five phenotype-associated publications detected via AMELIE for biallelic (D) DD and (E) DEE.

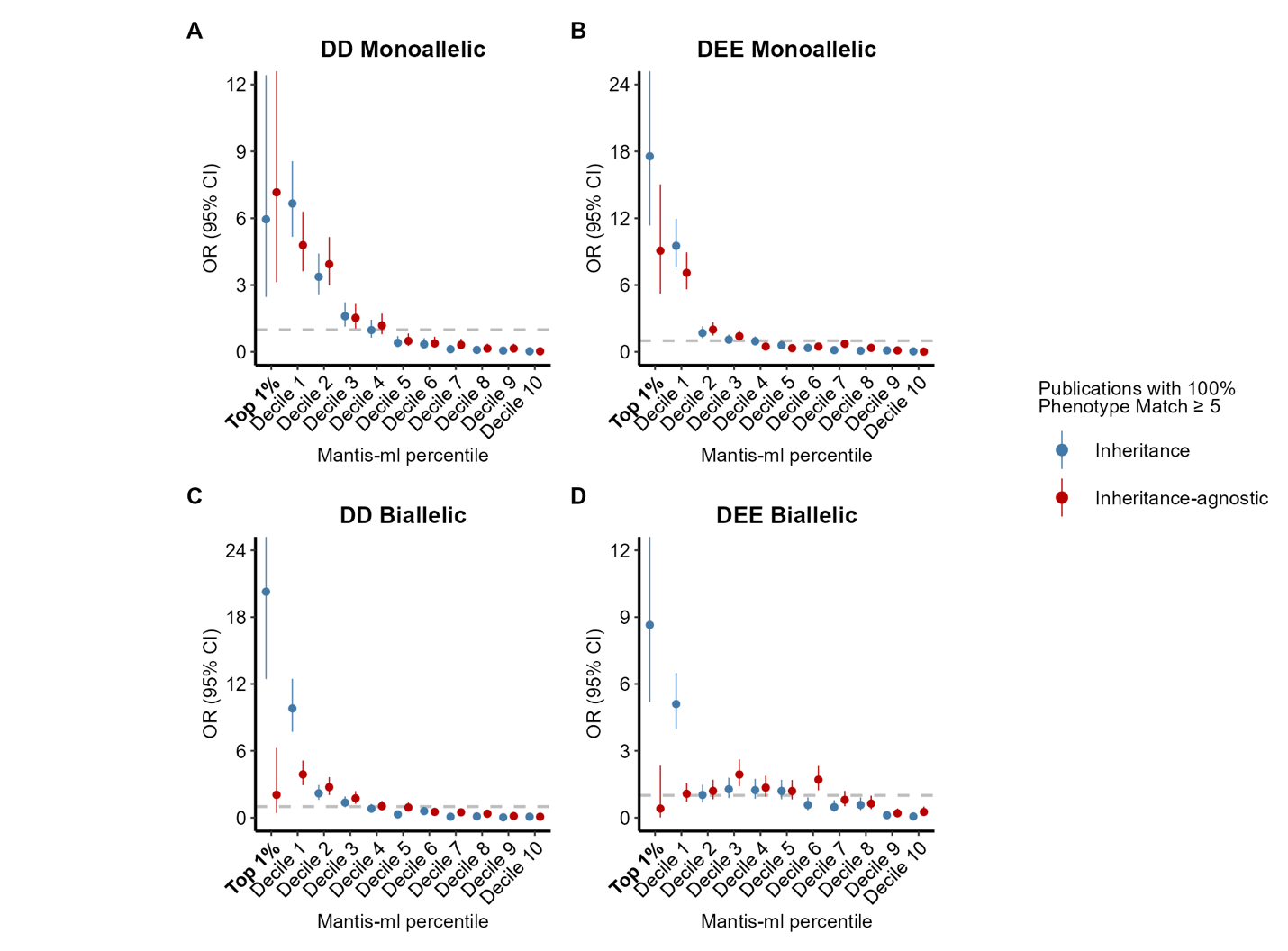

**Figure S11. Comparison of inheritance-informed and inheritance-agnostic model enrichment of genes with 100% phenotype match in published literature stratified by mantis-ml decile.** For each model, gene-phenotype match scores and related publications were ascribed to genes in an inheritance-specific manner (Methods). With seed genes removed, we selected genes with one or more publications assigned AMELIE gene-phenotype match scores of 100 and plotted the enrichment of the genes in the inheritance-stratified (blue) and inheritance-agnostic (red) prediction deciles.
